# Mining the CRBN Target Space Redefines Rules for Molecular Glue-induced Neosubstrate Recognition

**DOI:** 10.1101/2024.10.07.616933

**Authors:** Georg Petzold, Pablo Gainza, Stefano Annunziato, Ilaria Lamberto, Peter Trenh, Laura McAllister, Bradley Demarco, Laura Schwander, Richard D. Bunker, Mary Zlotosch, Rohitha SriRamaratnam, Samuel Gilberto, Gerasimos Langousis, Etienne J. Donckele, Chao Quan, Vaik Strande, Gian Marco De Donatis, Shanique B. Alabi, Jessica Alers, Michelle Matysik, Camille Staehly, Aurélie Dubois, Arnaud Osmont, Mackenzie Garskovas, David Lyon, Lars Wiedmer, Vladimiras Oleinikovas, Raphael Lieberherr, Nooreen T. Rubin, Daniel T. Lam, Nina Ilic Widlund, Andreas Ritzén, Ramon Miguel Caceres, Dominico Vigil, Jennifer Tsai, Owen Wallace, Marisa Peluso, Amine Sadok, Alison M. Paterson, Vladislav Zarayskiy, Bernhard Fasching, Debora Bonenfant, Markus Warmuth, John Castle, Sharon A. Townson

## Abstract

The CRL4^CRBN^ ubiquitin ligase is leveraged by molecular glue degraders, small molecules that reprogram ligase specificity to induce degradation of clinically relevant neosubstrate proteins. Known CRBN neosubstrates share a generalizable β-hairpin G-loop recognition motif, yet systematic exploration of the CRBN target landscape is still pending. Through computational mining of the human proteome using structure-based approaches, we predict over 1,400 CRBN-compatible β-hairpin G-loop proteins across diverse target classes, identify novel mechanisms of neosubstrate recognition through structurally differentiated helical motifs and molecular surface mimicry, and validate 22 representative neosubstrates with clinical implications. This work broadens the CRBN target space, redefines rules for neosubstrate recognition and establishes a platform for the elimination of challenging drug targets by repurposing CRL4^CRBN^ through next-generation molecular glue degraders.

## Main Text

Molecular glue degraders (MGDs) are an emergent class of small molecule drugs that induce proximity between ubiquitin ligases and target proteins causing target degradation through the cellular waste disposal system (*1*). Thalidomide analogs, particularly, are clinically effective MGDs that bind CRBN (*2*), the substrate receptor of the CRL4^CRBN^ ubiquitin ligase, and rewire substrate selectivity by creating a composite CRBN/drug neosurface that is competent in engaging so-called neosubstrate proteins (*3–5*). Neosubstrate recruitment is facilitated by drug-induced protein-protein interactions (PPIs) that drive cooperative complex formation with CRBN (*6–8*), which eliminates the need for pre-existing druggable pockets on the target and thus opens new avenues for drug discovery.

Previous studies on ternary complexes between CRBN, MGDs and known neosubstrates (e.g., CK1α, GSPT1, IKZF1, IKZF2, WIZ) have uncovered a common recognition motif that is shared across diverse structural domain types and protein classes (*6, 7, 9–12*), most considered undruggable in the past. The recognition motif is defined by a loop-like secondary structure element that is composed of a β-hairpin α-turn carrying a glycine residue at a key position of this loop (*6, 7*), a prevalent structural motif in the human proteome (*13*). These so-called β-hairpin G- loops define surface features on the target that create overall complementarity to the CRBN/MGD neosurface and enable productive protein-protein interactions with CRBN. Given the simplicity of the ternary complex interface, we expect that many more β-hairpin G-loop proteins will be accessible to this modality, and that target compatibility with CRBN may even go beyond the requirements defined by the β-hairpin G-loop paradigm.

### A proteome-wide map of the β-hairpin G-loop target space

To establish a map of the CRBN target space, we computationally mined the human proteome for motifs resembling *bona fide* β-hairpin G-loops from known CRBN neosubstrates. We chose the CK1α β-hairpin G-loop around glycine 40 (G40) as a query, selected from the CRBN:lenalidomide:CK1α ternary complex structure (*6*) (PDB id: 5FQD). We defined the CK1α β-hairpin G-loop by a stretch of eight amino acids (I35 to E42), comprising 5 residues N-terminal and 2 residues C-terminal of G40 (G-5 to G+2) (**Fig. 1A** and **B**). Together these residues preserve the geometry of the β-hairpin α-turn (*14*) through backbone-mediated intramolecular hydrogen bonds (G^…^G-4 and G-4^…^G+1; **Fig. 1B**), encompass the signature glycine (G0) and include two flanking residues, G-5 and G+2, whose amino acid side chains could engage in compound interactions (**Fig. 1B**). The query was matched against all human proteins available in two structural databases: the Protein Databank (PDB) and AlphaFold2 (AF2) models (*15*) of all annotated InterPro domains (*16*). We selected all structural motifs carrying a glycine residue that aligned to the query with a root mean square deviation (RMSD) smaller than 0.75 Å. We further triaged the identified hits by discarding matches sterically incompatible with CRBN after structural alignment to the β-hairpin G-loop of CK1α in PDB entry 5fqd (chain B and C; see methods). After removing protein domains primarily residing in the extracellular space, these stringent criteria identified 6,123 motifs distributed within a total of 1,424 human proteins (**Fig. 1C** and **Fig. S1A**). We recovered all previously reported β-hairpin G-loop targets (IKZF1, IKZF2, IKZF3, IKZF4, WIZ, ZFP91, ZNF692, GSPT1, ZMYM2, PDE6D, WEE1) (*3–5, 7, 9–12, 17–19*) along with a multitude of C_2_H_2_ zinc-finger (C_2_H_2_ ZF) proteins (**Fig. 1C** and **Fig. S1**). C_2_H_2_ ZF proteins are a well-established class in the β-hairpin G-loop target space (*9*) that encompasses a large majority of mined G-loops (5,160, 84% of all mined motifs) distributed over just 646 full-length proteins (**Fig. 1C** and **Fig. S1**). As ∼70% of all C_2_H_2_ ZF domains carry a β-hairpin G-loop (*9*) and often appear in domain clusters in the respective full-length proteins, individual C_2_H_2_ ZF proteins frequently contain multiple β-hairpin G-loop motifs. In contrast, the remaining 911 non-C_2_H_2_ β- hairpin G-loop motifs are distributed over 729 full-length proteins representing 96 different annotated protein classes and 272 domain types (**Fig. 1C**). 70% of these predictions are found in domains that lack obvious small molecule binding sites (*20*). Secondary structure assignment of 51 G-loops in 41 proteins was inconclusive due to deviation from canonical β-strand values, indicated as ‘Other’ in **Fig. 1C**.

**Fig. 1.**
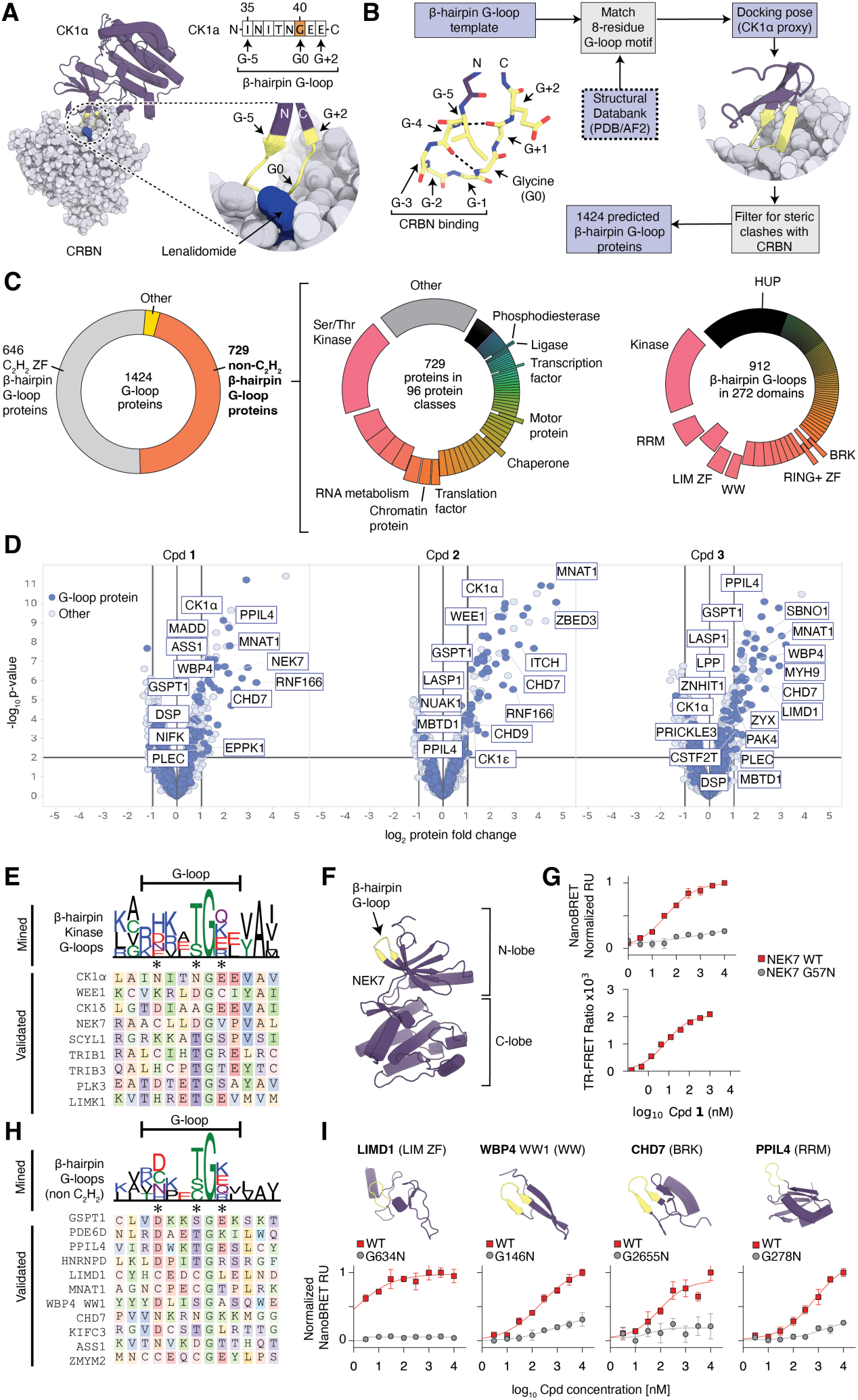
Exploration and validation of the β-hairpin G-loop target space. (**A**) CK1α:Lenalidomide:CRBN ternary complex (PDB id 5FQD) and close up of the CK1α β-hairpin G-loop degron (yellow). (**B**) Definition of the β-hairpin G-loop motif and schematic of the computational workflow. (**C**) Total distribution of β-hairpin G-loops across the proteome (left). Distribution of non-C_2_H_2_ β- hairpin G-loops across protein classes (center) and structural domains (right). Validated protein and domain classes are highlighted. (**D**) Volcano plots of TurboID experiments in CAL51 cells, 10 μM compound treatment for 3h. Proteins with predicted β-hairpin G-loops in non-C_2_H_2_ domains are highlighted. (**E**) Sequence logo plot of mined β-hairpin G-loops across kinase domains (top) and sequence alignment of β-hairpin G-loops in kinases validated in NanoBRET and TR-FRET (bottom). Asterisks highlight motif positions with amino acid preferences. (**F**) Structure of NEK7 kinase domain (PDB id 5DE2). (**G**) Ternary complex formation between CRBN and wild-type or mutant NEK7 validated in NanoBRET (top) and TR-FRET (bottom). (**H**) Sequence logo plot of mined non-C_2_H_2_ β-hairpin G-loops (top) and sequence alignment of validated non-C_2_H_2_, non-kinase β-hairpin G-loops (bottom). Asterisks highlight motif positions with amino acid preferences. (**I**) Ternary complex formation between CRBN and β-hairpin G- loop proteins representing novel protein domain classes.

Having established a computational map of the CRBN-compatible β-hairpin G-loop proteome, we next sought to validate compound-dependent CRBN recruitment of predicted targets using proximity-ligation experiments (*21*). For this we ectopically expressed a TurboID-CRBN construct in CAL51 cells to enable biotinylation of proteins that are proximal to CRBN in the presence of a compound. Proximity-labeled proteins were affinity-purified and analyzed by mass spectrometry using DMSO treated control cells as a reference. We profiled three compounds in these experiments: the previously reported FPFT-2216 (Cpd. **1**, (*22*)), a compound selected from our internal compound library (MRT-10350, Cpd. **2**), and CC-92480 (Cpd **3**, (*23*)) (**Fig. S2**). Proximity-labeling resulted in enrichment of numerous proteins in the presence of each compound (**Fig. S3A**), including the previously identified G-loop targets CK1α, WEE1 and GSPT1 (**Fig. 1D**), as well as many C_2_H_2_ ZF proteins **(Fig. S3B**). Apart from the well characterized targets, these experiments also identified novel β-hairpin G-loop proteins, including NEK7, MNAT1, PPIL4 and CHD7 (**Fig. 1D**).

Although our proximity-ligation experiments suggest a broad target range, indirect biotinylation of proteins interacting with *bona fide* G-loop targets can occur, as we predict for the FAM83 family of proteins through CK1α (*24*), and CDK7/CCNH through ΜΝΑΤ1(*25*) (**Fig. S3A**).Thus, we further validated direct target engagement and its dependence on the predicted G-loops using NanoBRET and TR-FRET. We initially focused these efforts on protein kinases, a class in which we mined 157 β-hairpin G-loops (**Fig. 1C**) located in the kinase N-lobe, in a position homologous to the G-loop of CK1α. We selected kinases observed in our proximity-ligation experiments (CK1α, WEE1, NEK7, SCYL1), and included additional ones that we predicted to carry a G-loop in the N-lobe, structurally diverge from the CK1α template by an RMSD between 0.22Å and 0.46Å (see methods), and feature potential amino acid preferences that emerge among the 157 kinase G- loops (CAMK2D, CK1δ, LIMK1, PLK3, STK4, TRIB1, TRIB3) (**Fig. 1E**). Using Cpd. **1-3** (**Fig. S2**), we observed ternary complex formation with CRBN for 9 of these kinases in both NanoBRET and TR-FRET (**Fig. 1G, Fig. S4** and **Fig. S5**), whereas CAMK2D and STK4 did not validate under these conditions. To demonstrate G-loop mediated recruitment to CRBN, we used the G-loop glycine to asparagine mutation that is predicted to result in clashes with the CRBN/MGD interface and was shown to prevent binding of other G-loop targets (*6*). Glycine to asparagine mutations within the identified targets ablated the NanoBRET signal (**Fig.1G, I** and **Fig. S4**), confirming G- loop-dependent CRBN engagement.

To further establish the generalizability of β-hairpin G-loops as a CRBN recognition motif, we next explored protein classes and domain types that have not been linked to this modality before. We selected a set of proteins with β-hairpin G-loop predictions for orthogonal validation in NanoBRET (PPIL4, MNAT1, WBP4, LIMD1, CHD7, ASS1, HNRNPD, RELB, GTF2B, MAD2L1, KIFC3; **Fig. 1H** and **Fig. S1**), some of which appear in our proximity-ligation experiments (**Fig. 1D**). These β-hairpin G-loops cover an RMSD range between 0.23Å and 0.66Å and are located in RRM, WW, BRK, LIM-ZF, RING ZF, kinesin or HUP domains (**Fig. S1** and **Fig. S4**).These proteins cover diverse functions ranging from chaperoning (PPIL4 (*26*)), splicing (PPIL4 (*27*), HNRNPD (*28*), WBP4 (*29*)), chromatin remodeling (CHD7 (*30*)) and transcriptional regulation (LIMD1 (*31*), MNAT1 (*25*)) to cellular trafficking (KIFC3 (*32*)) and metabolism (ASS1 (*33*)). Ternary complex formation for eight of these targets (PPIL4, MNAT1, WBP4 WW1 domain, LIMD1, CHD7, ASS1, HNRNPD, KIFC3) was validated in NanoBRET using Cpd. **1-3** (**Fig. 1I** and **Fig. S4**), whereas binding of RELB, GTF2B, MAD2L1 and the WW2 domain of WBP4 did not confirm under these conditions. Mutation of the predicted G-loop glycine to asparagine ablated binding (**Fig. S4**), confirming G-loop-dependent interactions with CRBN.

Sequence alignment of the validated β-hairpin G-loops revealed no discernible primary sequence motifs except for the glycine residue (**Fig. 1E, H** and **Fig. S1**). This is consistent with previous observations that sequence-diverse β-hairpin G-loops are compatible with CRBN, and that recruitment mostly relies on interactions driven by the backbone of the loop (*7, 9*). Although we do not observe obvious sequence requirements, amino acid preferences seem to emerge at positions G-4, G-1 and G+1 among the predicted G-loops outside the C_2_H_2_ ZF class (**Fig. 1E, H** and **Fig. S1**). These amino acid preferences are featured in the validated β-hairpin G-loop proteins (**Fig. 1E, H** and **Fig. S1**), suggesting that others will be compatible with CRBN as well. Overall, our systematic mining approach corroborates the β-hairpin G-loop as a generalizable CRBN binding motif that is present in a multitude of different domain types and target classes, many of which could be targeted through this modality.

### CRBN engagement through helical G-loop motifs

Our initial data together with previous reports suggest that target recruitment is sequence-diverse and largely driven by three G-loop backbone carbonyls (G-1, G-2, G-3) forming key hydrogen bond interactions with CRBN residues N351, H357 and W400 (*6–9*). As these interactions could in principle be satisfied by ternary folds other than β-hairpins, we repeated the mining using a less stringent definition of the G-loop. An analysis of the buried surface area of the CK1α G-loop engaged with CRBN:lenalidomide showed that residues G-3, G-2, G-1, G and G+1 contribute the largest part of the interface (**Fig. 2A**). We therefore reduced the query window to these 5 residues and repeated the mining with this minimal G-loop definition (G-3 to G+1). Using an RMSD cutoff of 0.5 Å and after elimination of motifs sterically inaccessible to CRBN and those in extracellular domains, this approach resulted in an increased number of predicted G-loop proteins (1,633), recovering all validated CRBN neosubstrates (**Fig. S1**) and 95% of those identified with the more stringent, 8 residue G-loop definition (G-5 to G+2). Interestingly, this approach produced 217 G- loop matches within helical parts of 184 full-length proteins (**Fig. 2B**), suggesting the existence of a helical G-loop motif (**Fig. 2B**). The FKBP12-rapamycin binding (FRB) domain of mTOR, a highly validated clinical target (*34*), contained such a helical motif with an RMSD of just 0.33 Å to the minimal G-loop template. NanoBRET confirmed ternary complex formation between the mTOR-FRB domain, CRBN and Cpd. **1** (**Fig. 2C** and **Fig. S6**). mTOR-FRB recruitment was dependent on the glycine residue G2092, confirming the predicted helical G-loop of mTOR-FRB as a CRBN recognition motif.

**Fig. 2.**
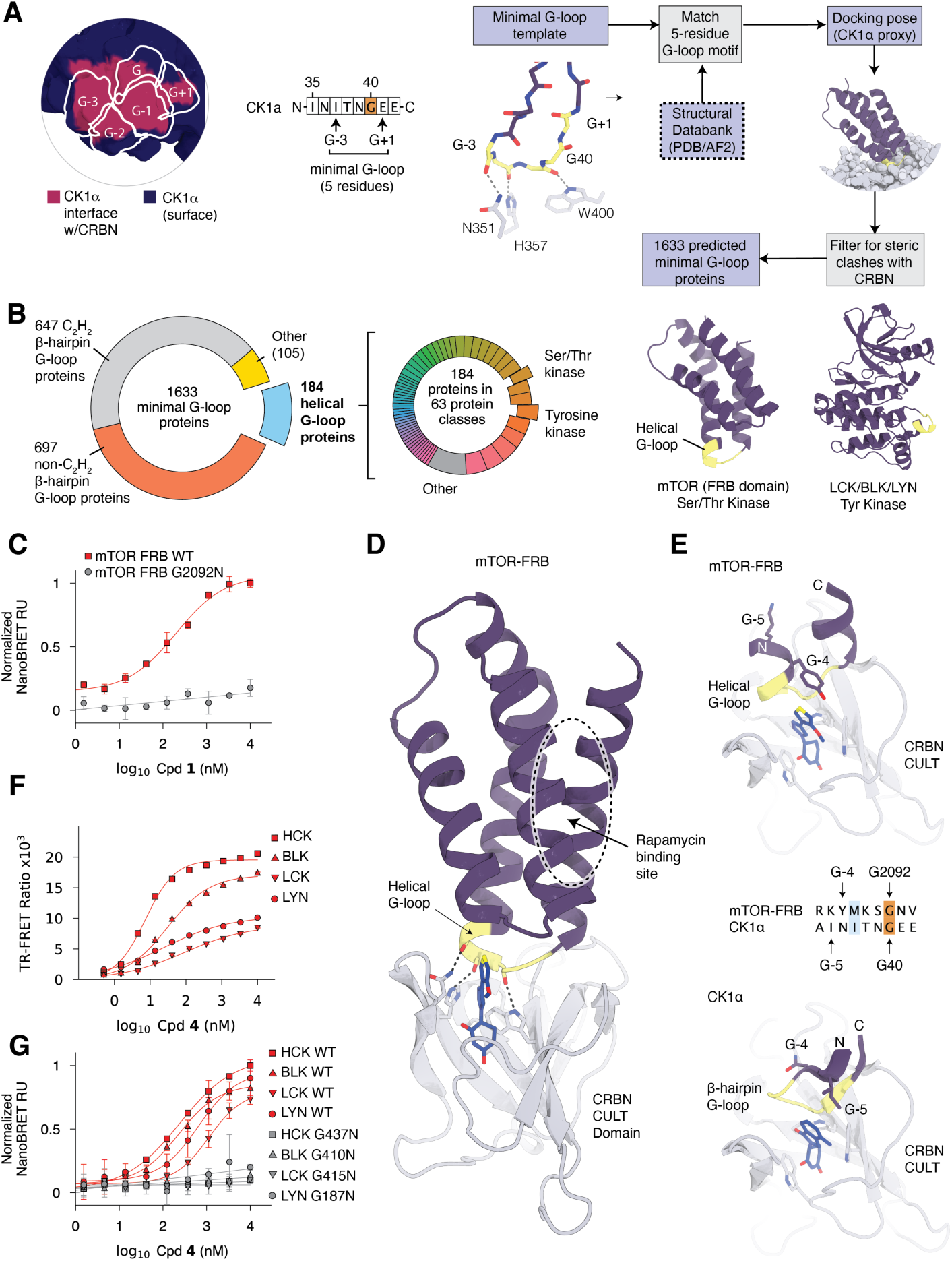
Discovery of structurally differentiated helical G-loop motifs. (**A**) CK1α β-hairpin G-loop surface with the region burried upon CRBN:Lenalidomide binding colored in red (left). Corresponding G-loop residues are highlighted and indicated in the CK1α G-loop sequence. Definition of the minimal G-loop and the corresponding computational workflow (right). (**B**) Distribution of the minimal G-loop target space across the proteome based on secondary structure identifies potential helical G-loop motifs (left). Distribution of helical G-loops across protein classes, with validated classes highlighted. Models of two domains predicted to carry a helical G-loop motifs. Helical G-loops colored in yellow (right). (**C**) NanoBRET validation of wild-type and mutant mTOR-FRB domain. (**D**) Ternary complex structure of CRBN, Cpd. **1** and mTOR-FRB. The CRBN CULT domain is shown in cartoon representation, other CRBN domains and DDB1 are omitted for clarity. (**E**) Comparison of the mTOR-FRB helical G-loop (top) and the CK1α β-hairpin G-loop (bottom) with differences at the compound interface highlighted. (**F**) TR-FRET validation of ternary complex formation between CRBN and kinases predicted to carry helical G-loop motifs. (**G**) G-loop mutations validate involvement of predicted helical G-loops of different kinases in ternary complex formation with CRBN in NanoBRET.

To investigate helical G-loop recruitment to CRBN at a molecular level, we determined a 2.95 Å crystal structure of the all-human ternary complex between DDB1^ΔB^-CRBN^Δ1-40^, Cpd. **1** and mTOR-FRB^2022-2114^. The helical G-loop at the apex of the four-helix bundle of mTOR-FRB engages CRBN through its C-terminal CULT domain (**Fig. 2D**), whereas the N-terminal LON domain of CRBN is dislodged and appears largely disordered in the crystal. The rapamycin binding site of mTOR-FRB is not in contact with CRBN or Cpd. **1**. Ternary complex formation is thus solely driven by the helical G-loop, which forms *bona fide* backbone interactions (G-3: M2089, G-2: K2090, G-1: S2091) with CRBN residues N351, H357 and W400. This constellation resembles recruitment of β-hairpin G-loops (**Fig. 2A**) and thus places the glycine and the G+1 residue (FRB G2092 and N2093) in the vicinity of the bound compound and CRBN residue V388, a hallmark of G-loop engagement by human CRBN (*35*). However, the backbone geometry of the G-4 residue (FRB Y2088) differs from that of β-hairpin G-loops due to changes in φ and ψ torsion angles of the helical fold. This exposes the side chain of G-4 (FRB Y2088) to the compound interface instead of the one of G-5 in the case of β-hairpin G-loops (**Fig. 2E**) thereby creating a more compact compound interface, which categorizes helical G-loops as a distinct CRBN recognition motif.

To further test the generalizability of helical G-loops, we selected the tyrosine kinases LCK, HCK, and LYN, whose predicted helical G-loops are located at the distal end of the αEF helix in the kinase C-lobe (**Fig. 2B** and **Fig. S7A**). The αEF helix is part of the kinase activation segment (*36*), which can adopt different conformations. Our mining approach only identified helical G-loops in LCK, HCK and LYN models representing the inactive state (**Fig. S7B**). Kinase models in an active conformation were found incompatible with CRBN due to steric clashes and were discarded by the algorithm (**Fig. S7C**). In NanoBRET, we observed recruitment for HCK with Cpd. **2** (**Fig. S7D**), whereas recruitment of LCK appeared weaker in these experiments. We therefore performed a TR-FRET library screen and identified Cpd. **4** as a potent inducer of ternary complex formation between CRBN and each of the three kinases (**Fig. 2F**), which could also be validated in NanoBRET (**Fig. 2G**). Mutation of the invariant G-loop glycine residue ablated ternary complex formation (**Fig. 2G**), validating our predictions. Interestingly, other SRC kinases, like BLK, lack structural information in the PDB, and the AF2 models represent a conformation that is incompatible with CRBN. Based on sequence conservation, however, the αEF helix of BLK could also carry a helical G-loop (**Fig. S6A**), which was confirmed by ternary complex formation with CRBN in the presence of Cpd. **4** (**Fig. 2F, G**). Taken together, our bioinformatic predictions establish that 1,633 proteins, a sizeable fraction of the human proteome, are in principle available for MGD-induced recruitment to CRBN via G-loop-like motifs. Whereas 1,424 carry CRBN-compatible β-hairpin G-loops, our analyses uncover a structurally differentiated helical G-loop motif present in 184 proteins that extends neosubstrate recognition by CRBN beyond β-hairpin α- turns.

### NEK7 binds CRBN through a differentiated PPI interface

Our computational mining approach highlights the prevalence of G-loops in the human proteome and establishes that distinct targets require differentiated chemistry for drug-induced proximity to CRBN (**Fig. S4**). Whereas CRBN proximity is an obvious requirement for CRL4^CRBN^-dependent polyubiquitination and degradation of a given target, proximity alone is not sufficient to result in degradation (*6, 9, 35*). Since our initial validation experiments focused solely on CRBN proximity, we next addressed whether degradation could be observed for some of the proximity-targets identified with Cpd. **1**. Whereas Cpd. **1** induced proximity of CK1α, NEK7, SCYL1, MNAT1, ASS1, CHD7, PPIL4, and several others to CRBN in recruitment-based readouts (**Fig. 1D**, **Fig. 2C** and **Fig. S3-S5**), only CK1α, PDE6D, and several C_2_H_2_ ZFs showed a >2-fold reduction of protein levels in global TMT proteomics (**Fig. 3A** and **Fig. S8A**). It has been shown that weak target recruitment can be converted into degradation through differentiated chemistry (*6, 9–12*). As we validated many G-loop targets through CRBN recruitment only, we wondered whether in some of these cases differentiated MGDs could in fact lead to degradation.

**Fig. 3.**
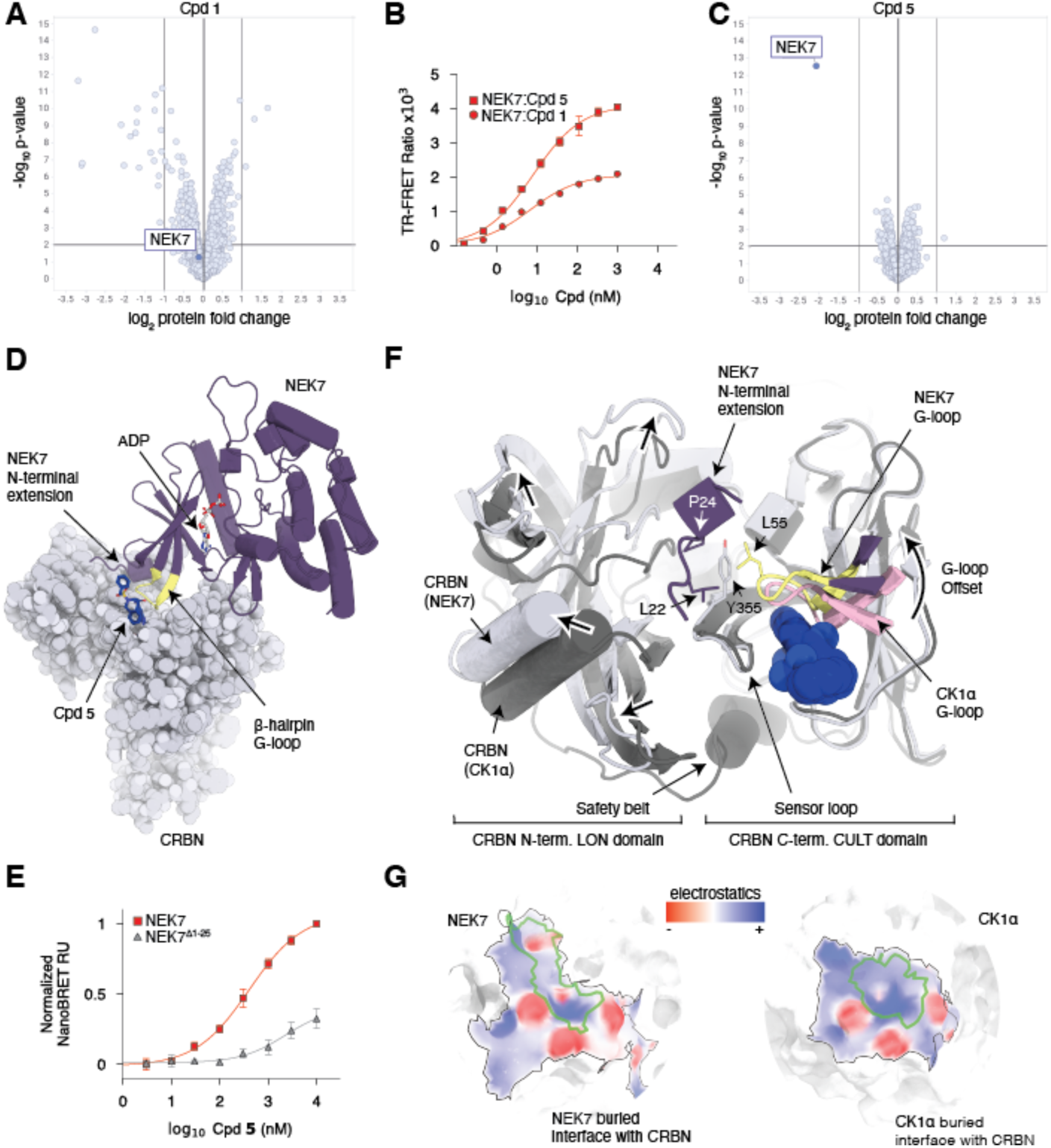
The novel β-hairpin G-loop target NEK7 uses a differentiated protein-protein interface with CRBN. (**A**) Global TMT proteomics in CAL51 cells, DMSO/MGD, 10 μM Cpd. **1**, 24 hours. (**B**) TR-FRET validation of compound-dependent NEK7 binding to CRBN. (**C**) Global TMT proteomics in CAL51 cells, DMSO/MGD, 10 μM of Cpd. **5** over 24 hours. (**D**) NEK7:CRBN:Cpd. **5** ternary complex structure. DDB1 omitted for clarity. (**E**) Normalized NanoBRET validation of CRBN engagement with either full length NEK7 or a NEK7 truncation lacking the first 25 residues at the N-terminus (NEK7^Δ1-25^). (**F**) Superposition of the NEK7:CRBN and CK1α:CRBN (PDB id: 5fqd) crystal structures using the CRBN CULT domain as a reference for alignment. Coloring: light gray/magenta (CRBN and NEK7 from NEK7:CRBN structure), dark gray/light pink (CRBN and CK1α from CK1α:CRBN structure). (**G**) Comparison of the electrostatics of the buried surface areas of NEK7 and CK1α when bound to CRBN in the presence of compound. Surface area contributed by the compound indicated by a green contour. Left: NEK7 buried interface in the DDB1^ΔB^-CRBN^Δ1-40^:Cpd. **5:**NEK7 crystal structure. Right: CK1α in the DDB1:CRBN:Lenalidomide: CK1α structure (PDB id: 5fqd).

To test this hypothesis, we turned our attention to NEK7, a NIMA-related serine/threonine kinase with therapeutic relevance in autoinflammatory diseases (*37*). NEK7 carries a β-hairpin G-loop in the N-lobe of its kinase domain with striking similarities to CK1α (**Fig. 1F** and **Fig. S1**), forms ternary complexes with CRBN and Cpd. **1** in TR-FRET, NanoBRET and TurboID proximity-ligation (**Fig.1D, G** and **Fig. S4-S5**), yet NEK7 steady-state levels remained unaffected in CAL51 cells over 24h using 10μM of Cpd. **1** (**Fig. 3A**). We therefore decided to screen our internal compound library for complex formation between NEK7 and CRBN in TR-FRET (**Fig. 3B**). This screen identified Cpd. **5**, which induced ternary complex formation with CRBN in both TR-FRET and NanoBRET, produced comparable EC_50_ values, but resulted in overall higher signal intensities compared to Cpd. **1** (**Fig. 3B** and **Fig. S8B**). Interestingly, treatment of CAL51 cells with Cpd. **5** led to potent NEK7 degradation (**Fig. 3C** and **Fig. S8C**), confirming that non-productive CRBN recruitment can be enhanced by MGDs to induce degradation of a given target (*6, 9–12*).

To dissect the ternary complex interface, we determined a 3.25Å crystal structure of an all-human ternary complex composed of DDB1^ΔB^-CRBN^Δ1-40^, Cpd. **5** and full-length NEK7 (**Fig. 3D**). Cpd. **5** binds CRBN through its dihydrouracil moiety buried in the tri-tryptophan cage, whereas the imidazopyridine ring and the branched tail are displayed on the CRBN surface available for interactions with NEK7. NEK7 binds the composite CRBN/Cpd. **5** interface through its predicted β-hairpin G-loop centered around G57 in the vicinity of the imidazopyridine ring, which facilitates hydrogen bond formation with CRBN residues N351, H357 and W400 (**Fig. S8D**). The G-loop interactions place the G-5 residue of NEK7, A52, proximal to the amide linker of the compound which allows for steric accommodation of NEK7 without forming obvious direct interactions with Cpd. **5**. Instead, the benzylsulfonamide moiety approaches more distal NEK7 residues and is in hydrogen bond distance to R35 and E37 that are located in a neighboring β-strand of the N-lobe (**Fig. S8E**). In addition, the structure revealed an N-terminal extension of NEK7 including L22, P24, and D25 that seem to extend the PPI with CRBN beyond the G-loop interactions (**Fig. 3D** and **Fig. S8F**). Deletion of these N-terminal residues in NEK7 (NEK7^Δ1-25^) reduces the NanoBRET signal (**Fig. 3E**), demonstrating that the extended PPI interface with CRBN is critical for ternary complex formation.

Comparing the NEK7 ternary complex to that of CK1α revealed crucial differences at the CRBN interface (**Fig. 3F** and **Fig. S8G-M**). The NEK7 G-loop is offset by 1.8Å at the position of the glycine (**Fig. 3F**) which affects the overall pose of the NEK7 kinase domain on CRBN (**Fig. S8G**). This offset is accompanied by a compound shift that otherwise would clash with the β-hairpin G- loop pose of CK1α (**Fig. S8H, I**), a feature that likely contributes to the selectivity profile of Cpd. **5** (**Fig. 3C**). Furthermore, compared to CK1α, NEK7 binding stabilizes CRBN in a partially domain-opened conformation (**Fig. S8J**). The G-loop offset and this CRBN conformation are unique features of the NEK7 ternary complex. Both possibly are consequences of structural rearrangements required to accommodate NEK7 while preventing steric clashes with closed CRBN observed in our docking model (**Fig. S8K-M**). These differences in the ternary complex interface illustrate the versatility of CRBN to accommodate different target surfaces. Comparing the footprints of NEK7 and CK1α on CRBN in the presence of their respective compounds shows substantial differences in electrostatics and surface area coverage, which further highlights the diverging interface requirements of both targets (**Fig. 3G**). Taken together, despite obvious similarities between CK1α and NEK7, both being serine/threonine-kinases, carrying β-hairpin G- loops in homologous positions of their kinase N-lobes and being recruited to CRBN by Cpd. **1**, ternary complex formation relies on markedly different protein contacts. This suggests that interactions between CRBN and its targets are more malleable than previously anticipated.

### VAV1 engages CRBN through molecular surface mimicry of a known degron

Given the emerging versatility of CRBN to engage targets with diverse interface requirements (*38, 39*), we reasoned that other structural arrangements may exist that further depart from the G-loop paradigm, yet satisfy key interactions with CRBN. As PPIs are driven by complementary surface properties, we hypothesized that G-loop-independent CRBN binding could be achieved through molecular surface mimicry of known target interfaces. To explore this hypothesis, we modified our surface matching algorithms (*40, 41*) to detect surface similarity to known CRBN degrons (**Fig. 4A** and **Fig. S9**). We used these updated algorithms to screen clinically relevant and undruggable targets lacking G-loop predictions including the VAV1 protein (*42*). VAV1 is a key mediator of B- and T-cell receptor signaling (*43*) that acts as a guanine-nucleotide exchange factor (GEF) for Rho family GTPases (*44*). VAV1 contains seven folded domains (**Fig. S9A**), none of which is predicted to carry a G-loop. We computationally screened all seven VAV1 domains for surface similarities to the ternary complex interfaces of CK1α (PDB id 5FQD) and GSPT1 (PDB id 6XK9) with CRBN (**Fig. 4B**), both markedly different to each other in shape and electrostatics. This approach identified two candidate surface patches within two separate SH3 domains of the VAV1 protein. The first one, located in the N-terminal SH3 domain (SH3n), showed similarities to the CK1α β-hairpin G-loop and corresponds to an N-loop, a structural motif with high backbone similarity to G-loops in which the glycine is replaced by an asparagine (**Fig. S9B**). As the glycine to asparagine mutation is predicted to lead to unfavorable steric interactions with CRBN and the MGD (**Fig. S9B**), the identified N-loop is likely incompatible. The second surface patch, identified in the C-terminal SH3 domain of VAV1 (SH3c), shows striking similarities to the GSPT1 surface contacting CRBN in complex with CC-90009 (**Fig. 4B**) (*45*). Interestingly, molecular surface similarity of the second surface patch could be identified despite the lack of underlying structural homology to either GSPT1 or its β-hairpin G-loop motif (**Fig. 4B**).

**Fig. 4.**
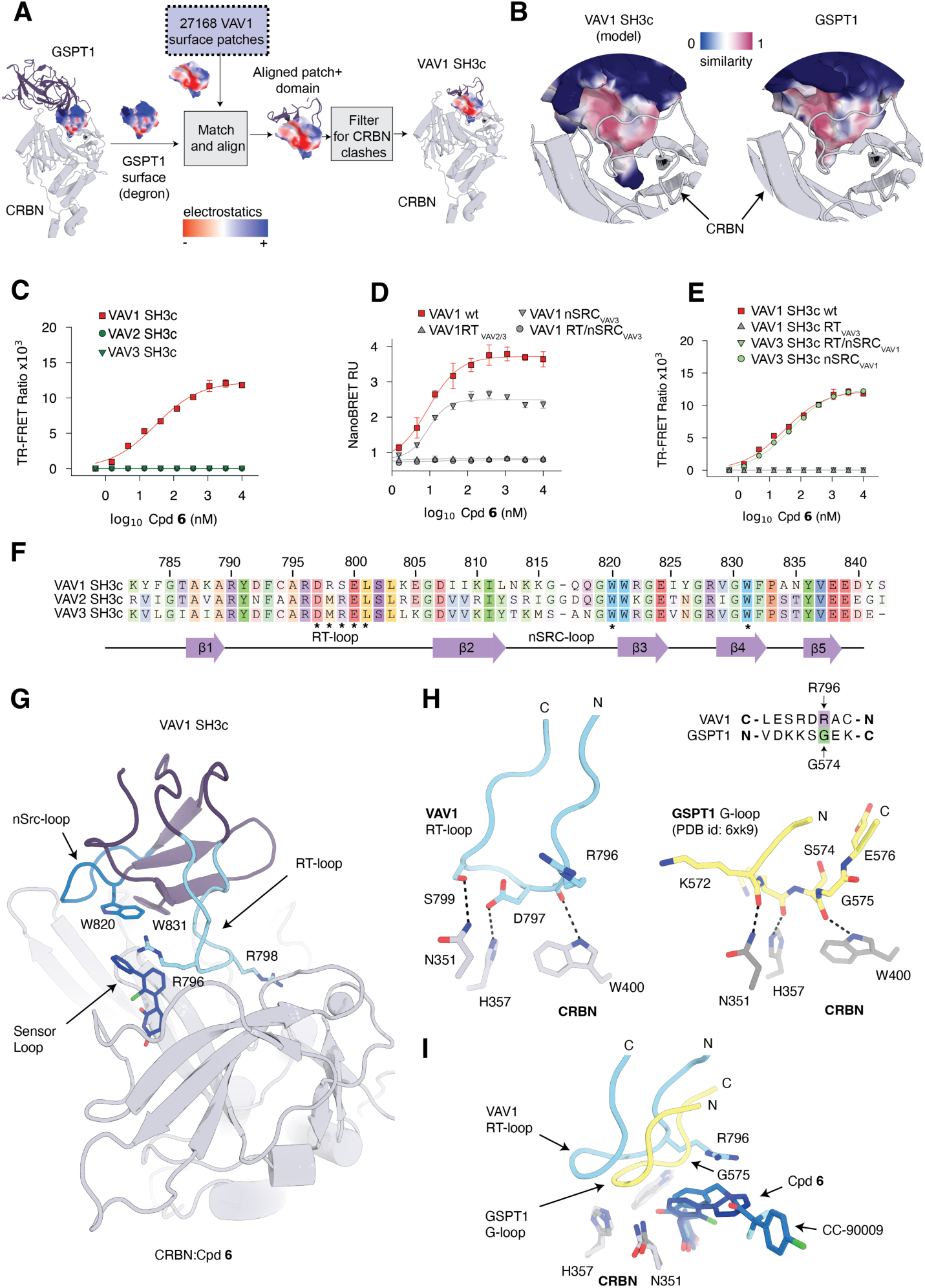
VAV1 binds CRBN through molecular surface mimicry of the GSPT1 degron. (**A**) Pipeline to match surface patches in VAV1 domains to known degron surfaces. (**B**) Left: Surface similarity between the predicted VAV1 SH3c pose and GSPT1 from the GSPT1:CRBN co-crystal structure (PDB id: 6xk9), with VAV1 SH3c colored according to surface similarity on a gradient from blue (no similarity) to red. Right: Coloring of the GSPT1 surface according to surface similarity to VAV1 SH3c in the model pose. (**C**) TR-FRET validation of CRBN recruitment of the SH3c domains of VAV1, VAV2, and VAV3. (**D**) NanoBRET validation of CRBN engagement with full-length wild-type and mutant VAV1. (**E**) Ternary complex formation in TR-FRET with wild-type VAV1 SH3c and chimeric mutant forms. (**F**) Sequence alignment of the SH3c domains of VAV1, VAV2 and VAV3. VAV1 residue numbering labeled above. Asterisks indicate residues contributing the VAV1 surface patch mimicking the GSPT1 degron. (**G**) VAV1:Cpd. **6**:CRBN ternary complex structure. DDB1 omitted for clarity. (**H**) Comparison of hydrogen bond interactions between CRBN and the VAV1 RT-loop and the GSPT1 G-loop (MGDs omitted for clarity). Left: VAV1 RT-loop; right: GSPT1 G-loop from GSPT1:CRBN:CC90009 structure (PDB id: 6xk9). (**I**) Superposition of VAV1 RT-loop and GSPT1 G-loop. Structural alignment was carried out via the CRBN CULT domain.

To probe compatibility of VAV1 SH3c with CRBN, we recombinantly expressed the VAV1 SH3c domain (residues 782-845) and performed a TR-FRET screen against our internal compound library. The screen identified Cpd. **6** as a potent inducer of ternary complex formation *in vitro*, and recruitment of full length VAV1 to CRBN through Cpd. **6** could be confirmed in cells using NanoBRET (**Fig. 4C-F**). The predicted surface patch in the VAV1 SH3c domain is created by two tryptophan residues, W820 and W831, and an apical region of the RT-loop (aa797-801; **Fig. 4F**). Whereas the tryptophan residues are conserved in the homologous SH3c domains of VAV2 and VAV3, their RT-loops diverge from the VAV1 sequence (aa798-799; **Fig. 4F**). Interestingly, neither VAV2-SH3c nor VAV3-SH3c led to ternary complex formation with CRBN in TR-FRET (**Fig. 4C**). Mutation of the RT-loop residues R798 and S799 in the SH3c domain of VAV1 to the respective VAV2/3 residues (M832 and R833 in VAV2) ablated binding in both NanoBRET and TR-FRET (**Fig. 4D, E**), whereas swapping nSRC-loop residues (aa812-818) still resulted in ternary complex formation (**Fig. 4D**). Changing RT-loop residues, but not nSRC residues, of VAV3 to the VAV1 sequence, reinstated CRBN binding (**Fig. 4E**), which establishes the RT-loop as a critical element for CRBN engagement.

To obtain structural insights into the binding mode of VAV1 SH3c, we solved a crystal structure of human DDB1^ΔB^-CRBN^Δ1-40^ in complex with Cpd. **6** and the VAV1 SH3c domain at 3.4 Å resolution (**Fig. 4G**). VAV1 SH3c engages the composite interface of CRBN and Cpd. **6** through the predicted surface patch in an orientation that deviates from the predicted pose by an interface RMSD of 2.0 Å (**Fig. S9C**). The RT-loop makes prime contacts with the G-loop binding site on CRBN, whereas the VAV1 residues W820 and W831 are positioned over the compound binding site and the sensor loop of CRBN. The nSRC-loop is located at the interface of the CRBN CULT and LON domains without forming obvious contacts in the structure. This constellation shows noticeable similarities to the pose of GSPT1 on CRBN despite striking differences in the ternary fold of both proteins (**Fig. S9D**). Like GSPT1, VAV1 forms hydrogen bond interactions with the three critical CRBN residues N351, H357 and W400 (**Fig. 4H**). Remarkably, however, unlike GSPT1, which engages N351 and H357 through backbone interactions of its G-loop, VAV1 engages these two CRBN residues through the side chains of S799 and D797 located in the RT-loop (**Fig. 4H**). The third CRBN residue, W400, forms a hydrogen bond with the backbone carbonyl of VAV1 residue R796 (**Fig. 4H**). R796 takes the analogous position of the G-loop glycine, with its side chain pointing towards Cpd. **6** (**Fig. 4H, I**). In addition, the RT-loop residue specific to the VAV1 sequence, R798, extends the PPI interface with CRBN through its elongated side chain (**Fig. 4G**). The remarkable differences in the hydrogen bond architecture between the GSPT1 G-loop and the VAV1 RT-loop result from reversed orientations of their N- and C-termini (**Fig. 4H, I**). Owing to resolution limits, data quality or conformational flexibility, we only observed electron density for the 2-chloro phenyl-glutarimide of Cpd. **6**, whereas the distal *N*- methyl pyrazole remained unresolved and was omitted from the final model.

Taken together, our structural analyses illustrate how VAV1 mimics hydrogen bond interactions normally observed between CRBN and G-loops and rationalizes key contributions of VAV1 residues in CRBN engagement. Furthermore, the VAV1 SH3c domain shows striking molecular surface mimicry to the GSPT1 interface with CRBN/CC-90009 in the absence of underlying structural homology. These findings redefine rules for neosubstrate recognition and suggest that other proteins lacking G-loops can engage CRBN through molecular surface complementarity.

## Discussion

Our systematic approaches to chart the CRBN target space identified 1,633 human proteins that we predict to be compatible with CRBN through accessible β-hairpin or helical G-loop motifs. Almost 1,000 of these lie outside the well characterized C_2_H_2_ zinc-finger class (*9*), represent diverse protein functions and domain types, including many that are currently perceived inaccessible to small molecule ligands. Experimental validation of 21 novel β-hairpin and helical G-loop proteins quadruples the number of confirmed non-C_2_H_2_ CRBN targets in the public domain, corroborates the sequence-diverse nature of the G-loop motif and establishes that different G-loop targets require differentiated chemistry for CRBN engagement. As only four compounds have been used to validate a representative set of 21 targets, our data strongly suggest that other members of the computationally mined G-loop proteome will be accessible to CRBN through differentiated chemistry.

Despite G-loops representing a universal structural motif that is characterized by its ternary fold and the identity of the glycine residue, target recruitment is eventually driven by topological and electrostatic features on the protein surface that create overall complementarity to the CRBN/MGD neosurface. Whereas previously discovered CRBN targets all share a β-hairpin G-loop, VAV1 binds CRBN in a G-loop-independent manner through molecular surface mimicry of the GSPT1 degron, a well-established CRBN-dependent G-loop neosubstrate. As VAV1 lacks detectable sequence similarity or structural homology to GSPT1, conserved primary, secondary and ternary elements are not a requirement to create CRBN compatible surface features. These findings highlight the extraordinary plasticity of a primary interaction site on CRBN (CRBN residues N351, H357, W400) that is compatible with the G-loop motif, but also with the differentiated binding mode of VAV1. This primary interaction site is a central element of a larger PPI hotspot on the CRBN surface, which has high predicted propensity to engage in protein-protein interactions (**Fig. S10A**). As observed for NEK7, VAV1 and other structurally characterized neosubstrates (*7, 9, 11, 38, 39, 46*), the PPI hotspot on CRBN provides opportunities for cooperativity through MGD-induced, target-specific interactions (**Fig. S10B**). Interestingly, the PPI hotspot spans two CRBN domains, the LON and the CULT domains, which are flexibly attached (*47*) and allow for conformational adjustments to maximize target complementarity, as exemplified by the NEK7 ternary complex structure (**Fig. 3F** and **Fig. S8**).

Altogether, our results offer opportunities for CRBN target space expansion within and beyond the β-hairpin G-loop paradigm. We demonstrate that neosubstrate complementarity will be a function of the CRBN PPI hotspot, the conformational flexibility on both sides of the ternary complex interface, as well as the ability to modulate central surface features through MGD diversification. Future computational mining approaches will model complementarity between molecular surfaces present in the human proteome and the vast diversity of possible CRBN/MGD neosurfaces for the discovery of novel CRBN targets with therapeutic relevance.

## Acknowledgments

We thank Xavi Lucas and Elisa Liardo for their assistance with chemical compound analysis. We thank Irina Cornaciu-Hoffmann and the entire team at ALPX, Grenoble for crystallization and data collection support. We are grateful to Ralph Tiedt for feedback and comments on the manuscript.

## Author contributions

PG, GP, RDB: conceptualization of G-loop mining including motif windows, and mimicry searches. PG, MC: software code. SA, SG, GD, RL: β-hairpin G-loop validations, NEK7 G-loop validation in NanoBRET. IL, MZ, GL, SBA, NIW, VZ: helical G- loop validations, VAV1 NanoBRET. LS, VS, AO, AD, GL, DL, DB: Mass spectrometry Turbo ID/global TMT; neosubstrate identification. BD, JA, MG, DV: TR-FRET experiments and library screens for LCK, NEK7, VAV1. BD, JT, PT: recombinant protein sourcing. RS, MM, CS, AS: NEK7 target. PT, SBA, JA, VO, BD, MP, AMP: VAV1 degron validation. LM, ED, AR, BF: MGD chemistry conceptualization. GP, PT, CQ, JT: Crystallography. RDB: Crystallography data processing lead. VO, LW: CADD support. GP, PG narrated and drafted the manuscript. GP, PG, JC, SAT reviewed and edited the manuscript. All authors analyzed data, reviewed, and commented on the manuscript. MW, JC, SAT: supervised research.

## Data and materials availability

All software code, intermediate files, and predictions will be deposited in zenodo.org. All software code will be available at: http://github.com/monterosatx/gloop_mining and http://github.com/monterosatx/molecular_surface_mining. Protein Data Bank (PDB) files will be deposited into the Research Collaboratory for Structural Bioinformatics (RCSB) PDB.

---

**Fig. S1.**
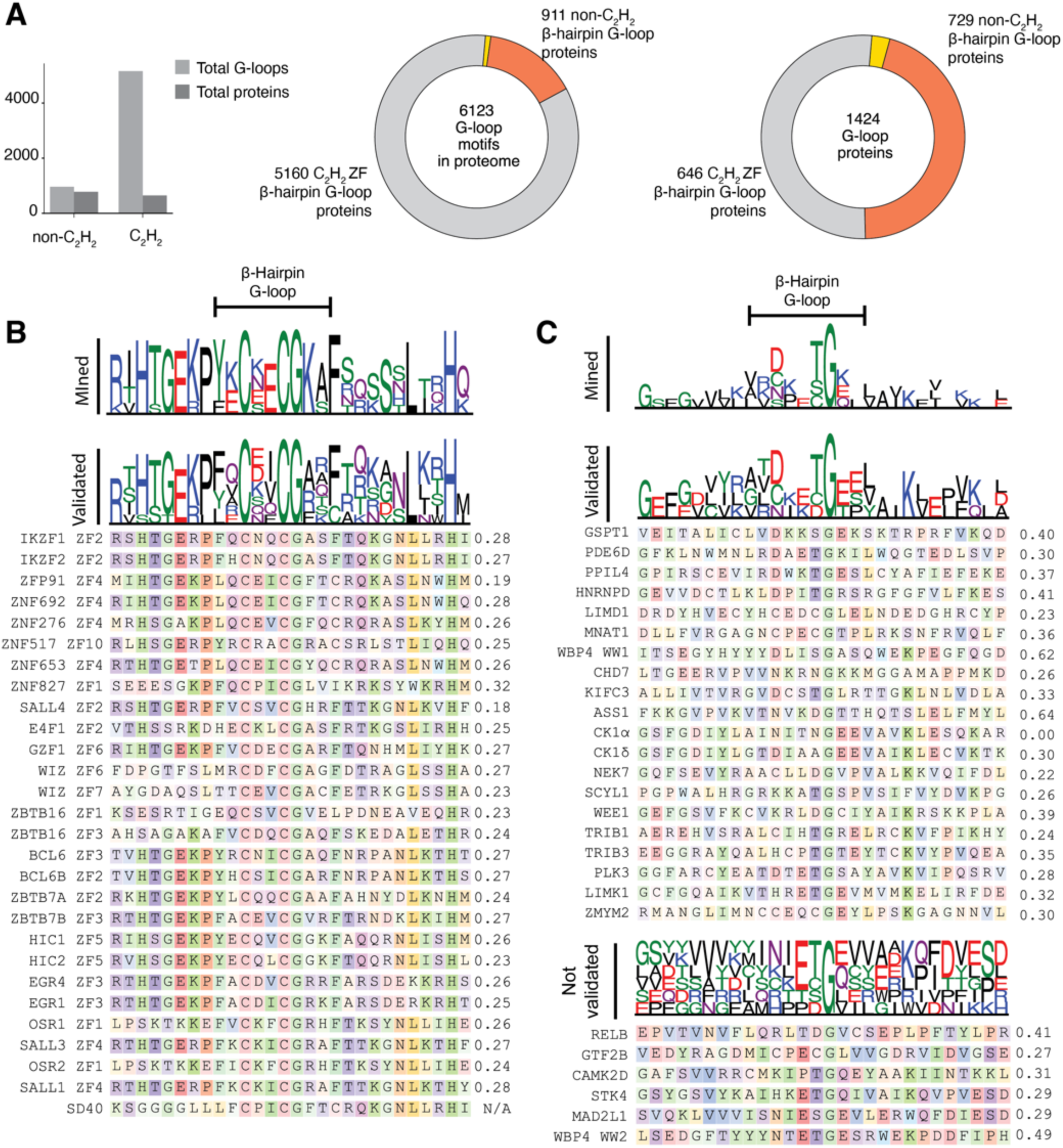
Sequence logo for Mined, Validated and β-hairpin G-loops that did not validate experimentally. (A) Left: Absolute number of matched motifs in either C_2_H_2_ ZFs or non-C_2_H_2_, and distribution in full length proteins. Mined motifs matching the β- hairpin G-loop query in the human. (B) Top: Sequence logoplot for C_2_H_2_ ZFs mined by our method. Middle: logoplot for C_2_H_2_ ZFs validated in the literature. Bottom: sequence alignment of C_2_H_2_ ZFs validated in the literature, spanning the range from G-14 to G+13. (C) Top: Sequence logo plot for mined β-hairpin (non-C_2_H_2_) G-loops in this work. Middle: sequence logo plot and sequence alignment of β-hairpin (non-C_2_H_2_) G-loops validated in this paper, in addition to previously validated GSPT1, PDE6D and ZMYM2. Bottom: Logo plot and sequence alignment of the G-loop region for β-hairpin (non-C_2_H_2_) G-loops that were tested under either Cpd 1, 2, or 3 in NanoBRET in this work but did not show a binding signal. The number next to each aligned sequence in all panels shows the backbone atom root mean square deviation between the predicted G-loop and the CK1α query G-loop in the mining.

**Fig. S2.**
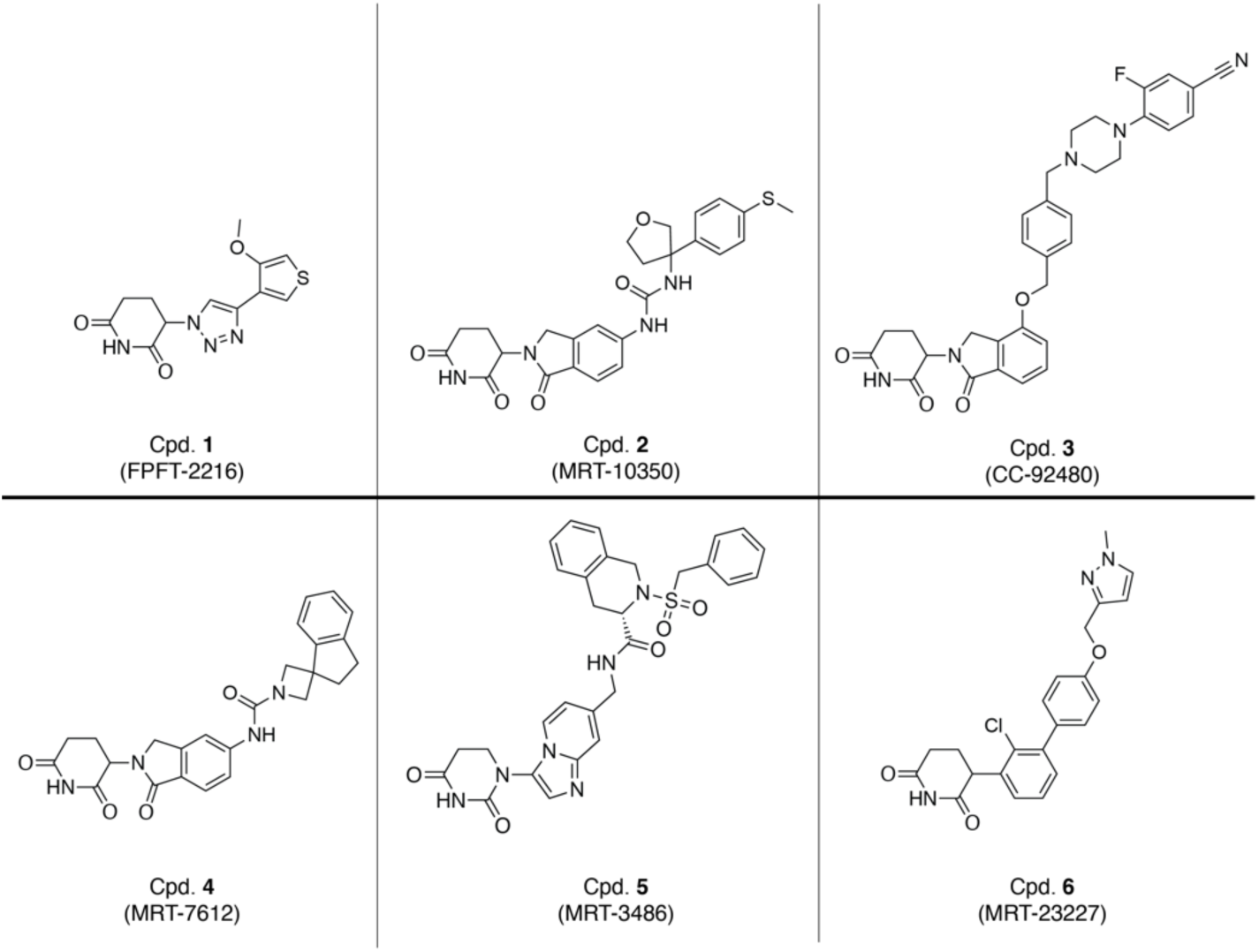
Structures of the chemical compounds used in this work.

**Fig. S3.**
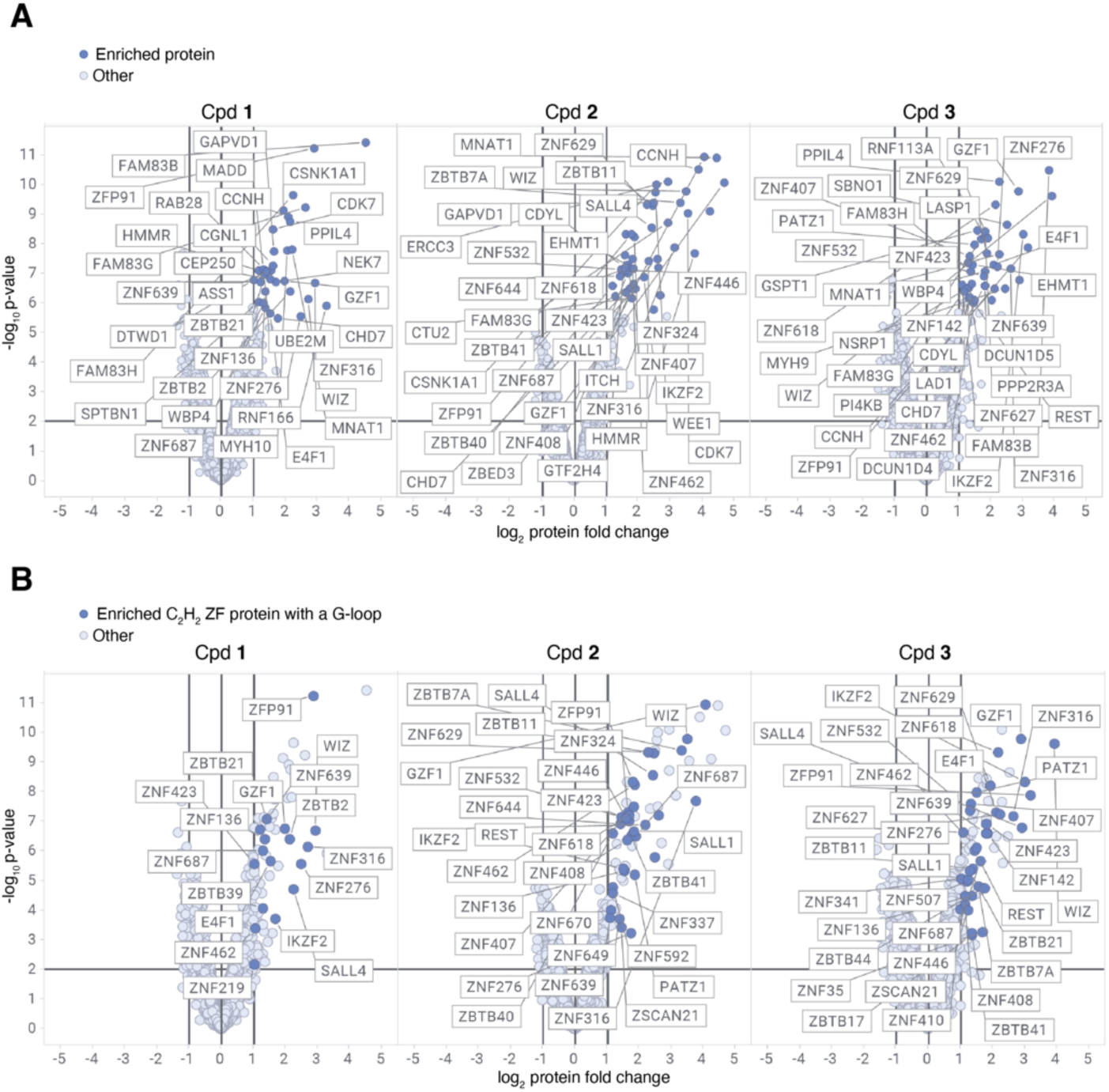
Proximity labelling of proteins in TurboID+proteomics. (A-B) Volcano plot of biotinylated protein enrichment based on mass spectrometry analysis of CAL51 cells exposed to either 10 μM Cpd or DMSO and harvested after 3 hours. (A) All enriched proteins (dark blue circles) are labelled. (B) Only enriched proteins with a predicted β-hairpin G-loop on a C_2_H_2_ ZF domain are labelled (dark blue circles).

**Fig. S4.**
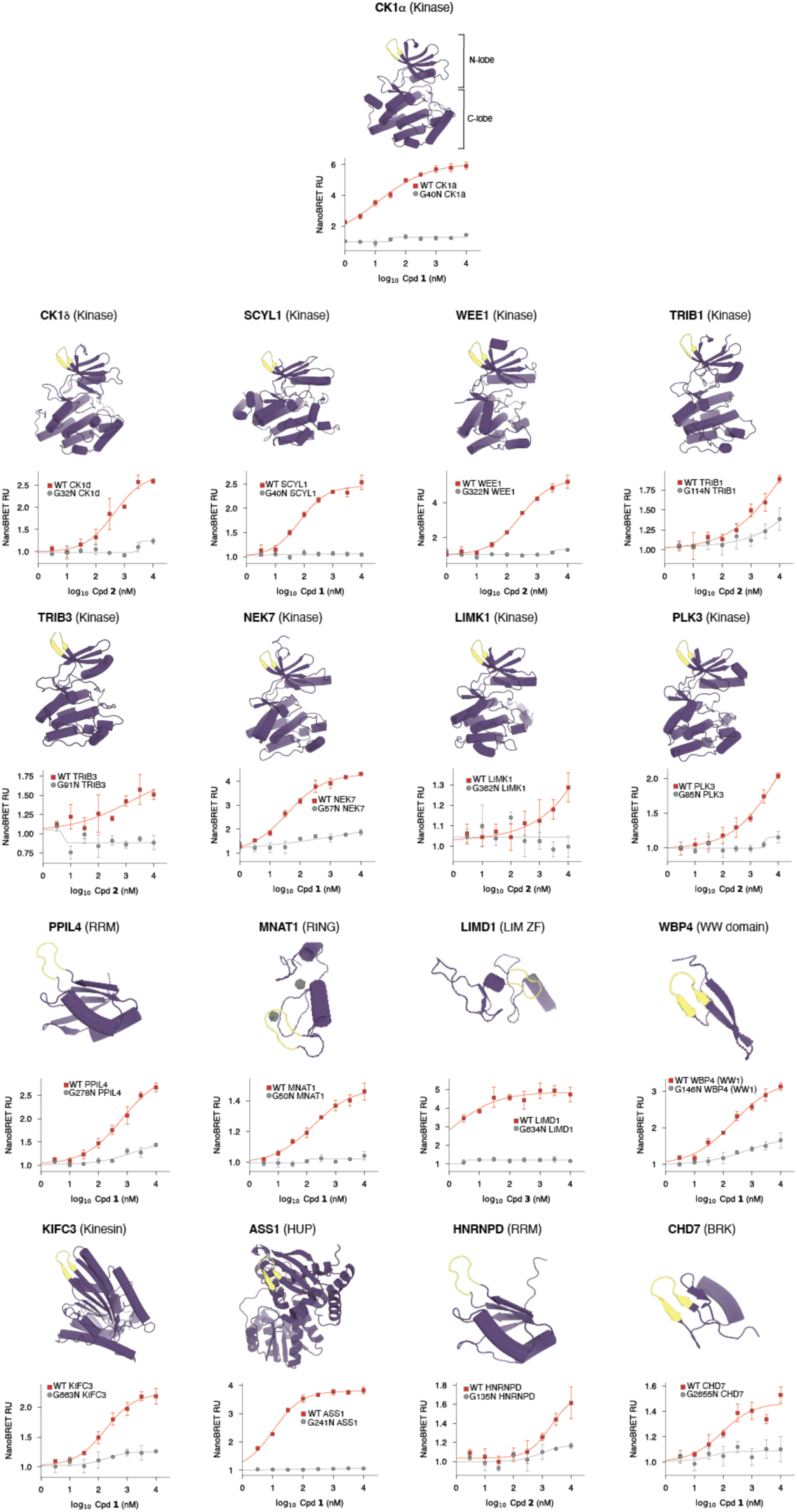
NanoBRET validation of β-hairpin G-loops across human domains. Each panel shows a rendering of a β-hairpin G- loop-containing domain validated in this work, with the protein name above the rendering (in bold) and the domain type in parentheses. Each plot below shows the NanoBRET validation of either WT binding of the domain or a mutation in the central glycine of the β-hairpin G-loop, under increasing concentrations of Cpd. The specific compound used for each NanoBRET experiment is indicated.

**Fig. S5.**
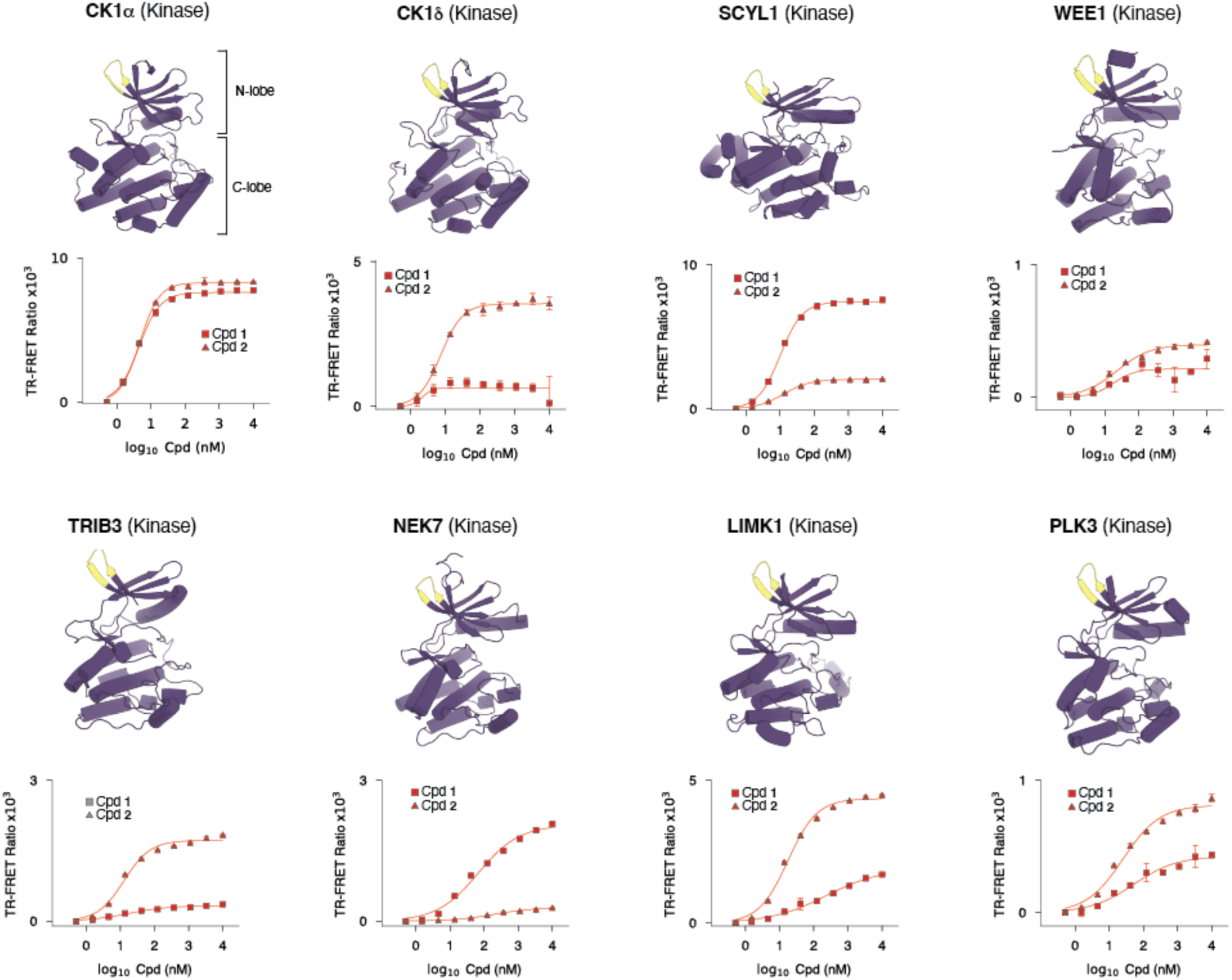
TR-FRET validation of validated kinases with Cpd. 1 and Cpd. 2. Each panel shows a rendering of a β-hairpin G- loop-containing kinase validated in this work, with the protein name above the rendering (in bold). The plots below show TR-FRET validation of the recombinant wild type kinase binding to CRBN:DDB1 under increasing concentration of either Cpd. **1** (red boxes) or Cpd. **2** (red triangles).

**Fig. S6.**
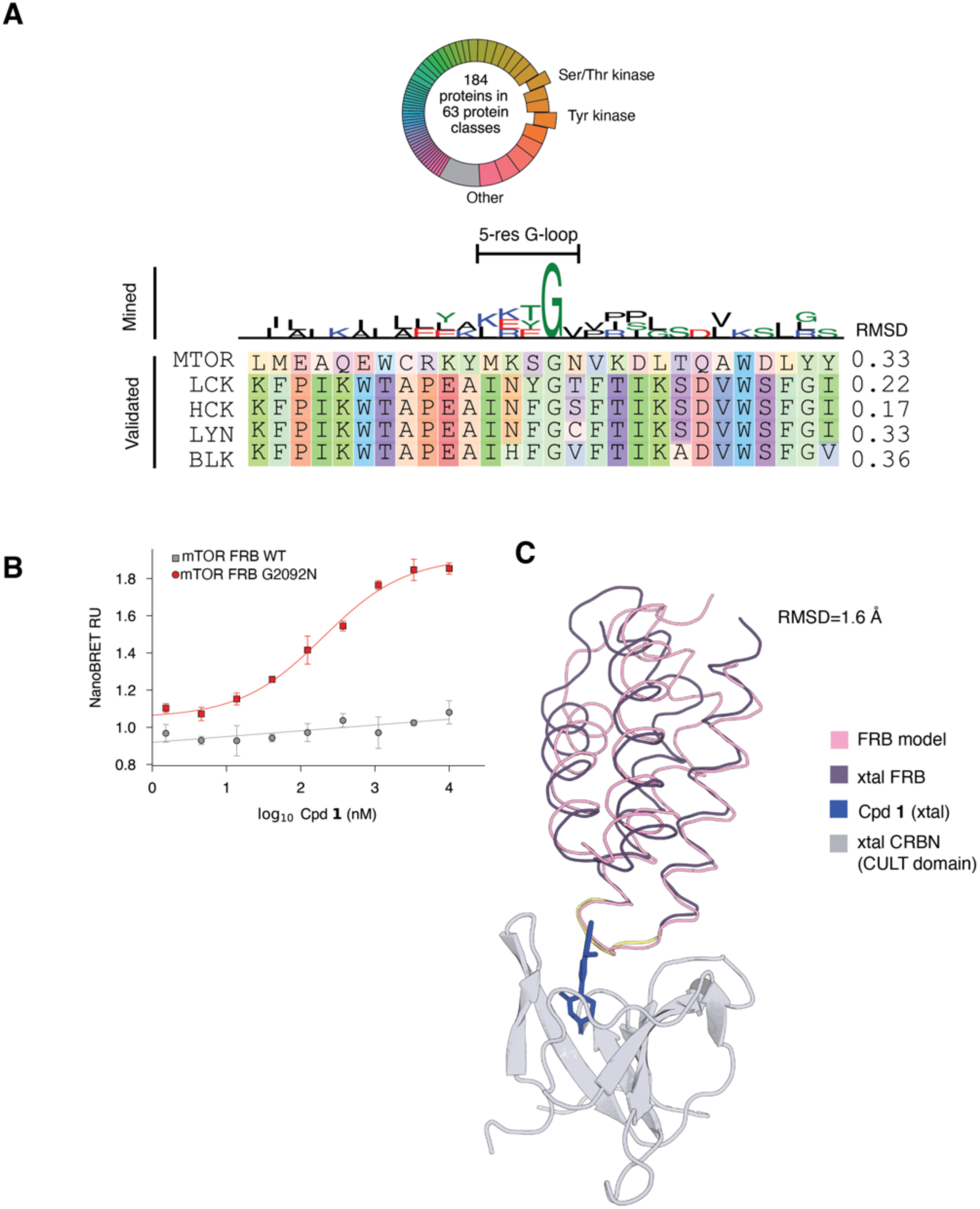
Sequence analysis of helical G-loops, and analysis of mTOR-FRB helical G-loop. (A) Above: Distribution of helical G-loops on protein domains. Below: Sequence logo plot of the mined G-loops, showing a region between 14 residues before and 13 after the central glycine, and sequence alignment of the validated helical G-loops in this paper. (B) NanoBRET validation of wildtype mTOR FRB and the mutant G2092N. (C) Comparison between predicted CRBN:FRB model and crystal structure of CRBN:FRB domain. The CULT domain of the predicted CRBN:FRB model was aligned to the CULT domain of the solved crystal (xtal) structure. Rendering shows the position of the FRB model (pink wires) compared to the FRB crystal.

**Fig. S7.**
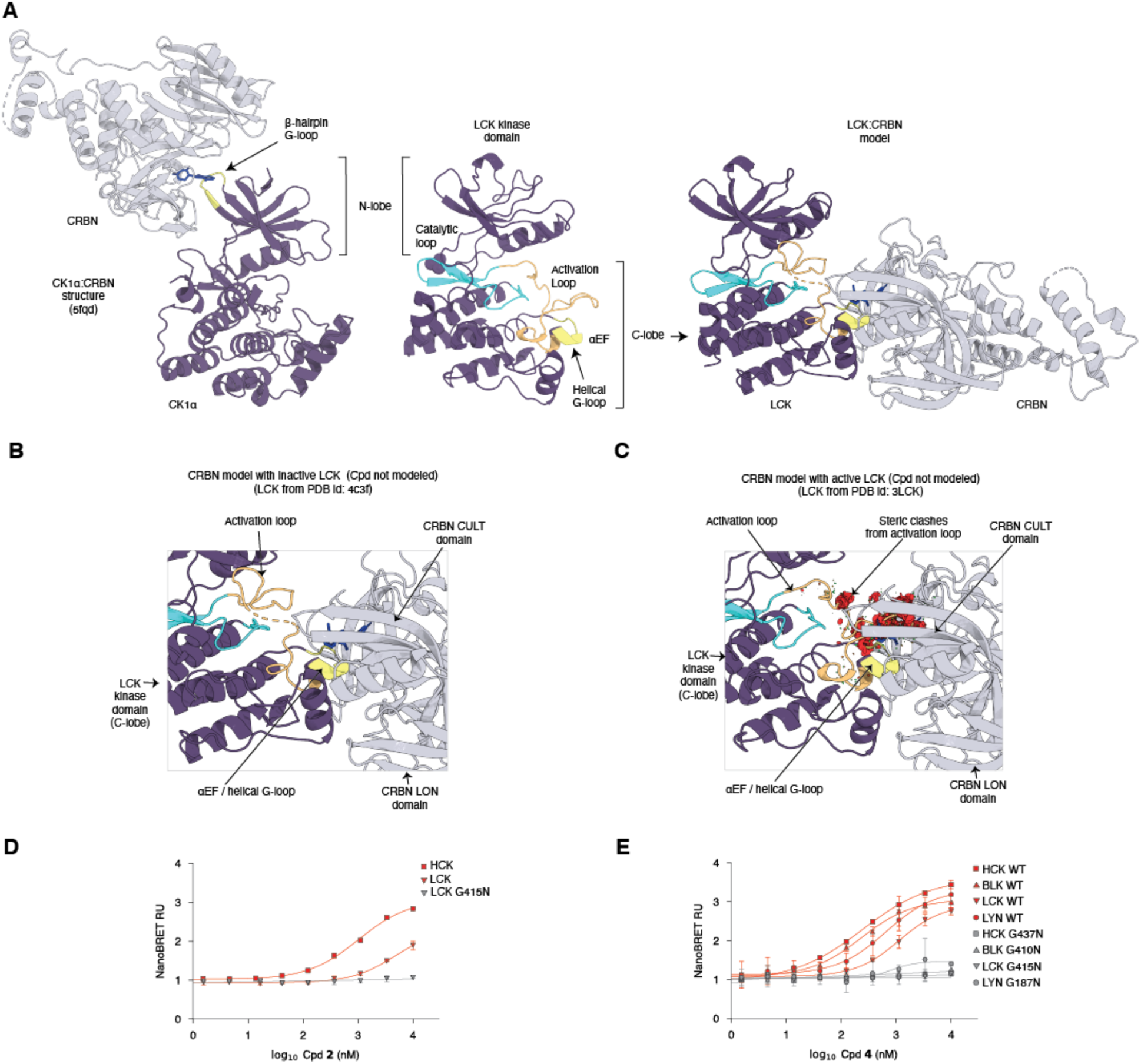
Modeling and additional validation of the helical G-loop in SRC kinases. (A) Kinase recruitment to CRBN. (left) CK1α recruitment to CRBN through β-hairpin G-loop in the kinase N-lobe. (center) Model of the LCK kinase domain. Activation segment comprising catalytic loop, activation loop and αEF helix are highlighted. (right) Model of LCK engaging CRBN through its helical G-loop. (B) Modeling of inactive conformation of LCK with missing activation loop residues. (C) Modeling of LCK active conformation with predicted helical G-loop in complex with CRBN, with clashes shown in red. (D) NanoBRET validation of Cpd. 2 binding to HCK, LCK and the predicted helical G-loop mutant G415N. (E) NanoBRET validation of SRC kinases and G-loop ablation mutants with Cpd. 4.

**Fig. S8.**
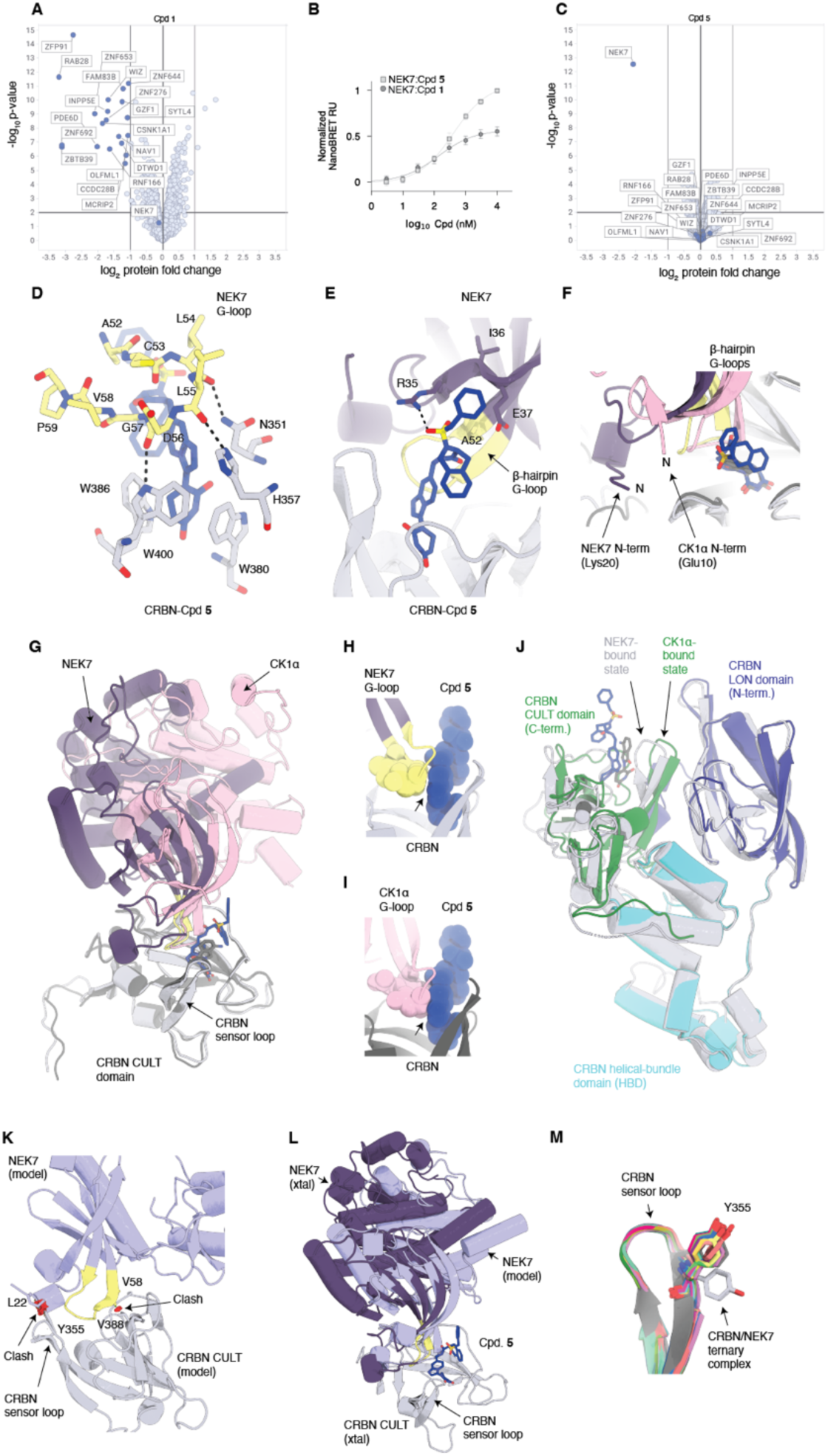
Proteomics, NanoBRET and structural characterization for NEK7 and Cpd. 5. (A) Degradation proteomics in CAL51 cells exposed to 10 μM of Cpd 1 over 24 hours. (B) NanoBRET validation of NEK7 binding to CRBN. (C) Global TMT proteomics in CAL51 cells exposed to DMSO/10 μM of Cpd 5 over 24 hours. (D) NEK7 G-loop binding in presence of Cpd. 5, with bona fide hydrogen bonds shown with a dotted line. (E) Cpd 6:NEK7 interactions. (F) Comparison between the NEK7 N-terminal extension (dark purple) and the homologous CK1α N-terminal region (pink). (G) Comparison between the NEK7 and CK1α binding pose (PDB id: 5fqd) after structural alignment of CRBN CULT domains. (H) Interaction of the NEK7 G-loop with Cpd. 5. (I) Modeling of CK1α with Cpd. 5. (J) Structural comparison of CRBN in the NEK7 and CK1α structures (PDB id: 5fqd) after CULT domain-based alignment. (K) Clashes between NEK7 and CRBN in the model using the CK1α:CRBN structure’s G-loop (PDB id: 5fqd) as a template for G-loop based alignment. (L) Binding pose comparison between NEK7 predicted model and NEK7 structure. (M) Comparison between sensor loop of 26 publicly available CRBN conformations from 15 non-PROTAC ternary complex structures compared to the sensor loop in the NEK7:CRBN structure.

**Fig. S9.**
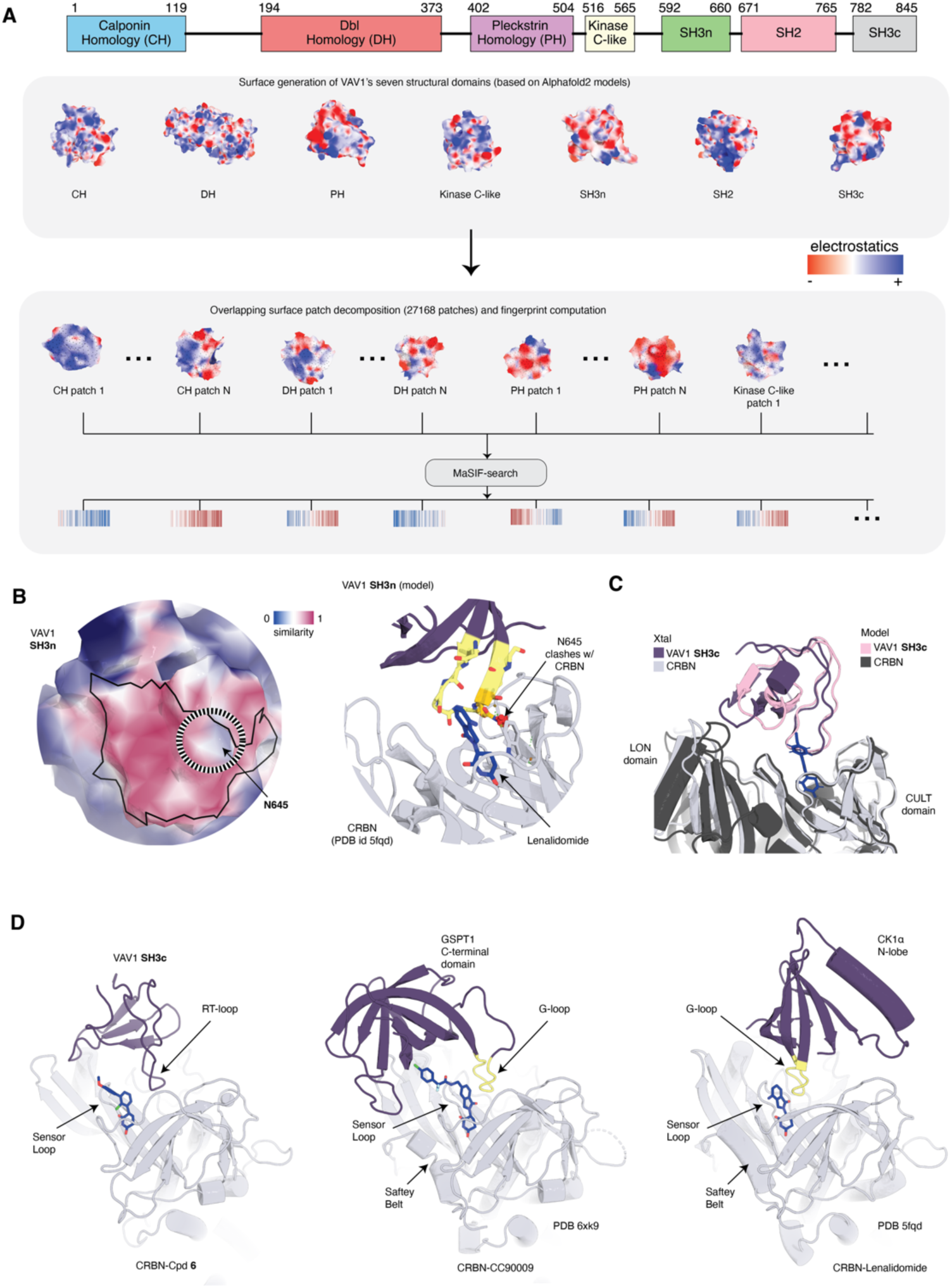
Surface modeling of VAV1 domains. (**A**) Top: Structural domain distribution in the VAV1 gene. Middle: molecular surface representation of the VAV1 structural domains, colored by the Poisson-Boltzmann electrostatics. Bottom: Decomposition of the VAV1 molecular surface into overlapping radial patches. (**B**) (left) Surface similarity of VAV1 SH3n to CK1α after a surface-based alignment with the N residue of the N-loop highlighted. (right) Structural view of the ternary SH3n CRBN (from PDB id: 5fqd) model after surface-based alignment. (**C**) Comparison between surface-based alignment model of VAV1 SH3c and crystal structure of the ternary VAV1 SH3c:Cpd. 6:CRBN complex. (**D**) Comparison of the binding modes of VAV1 SH3c, GSPT1 (PDB id: 6xk9) and CK1α (PDB id: 5fqd).

**Fig. S10.**
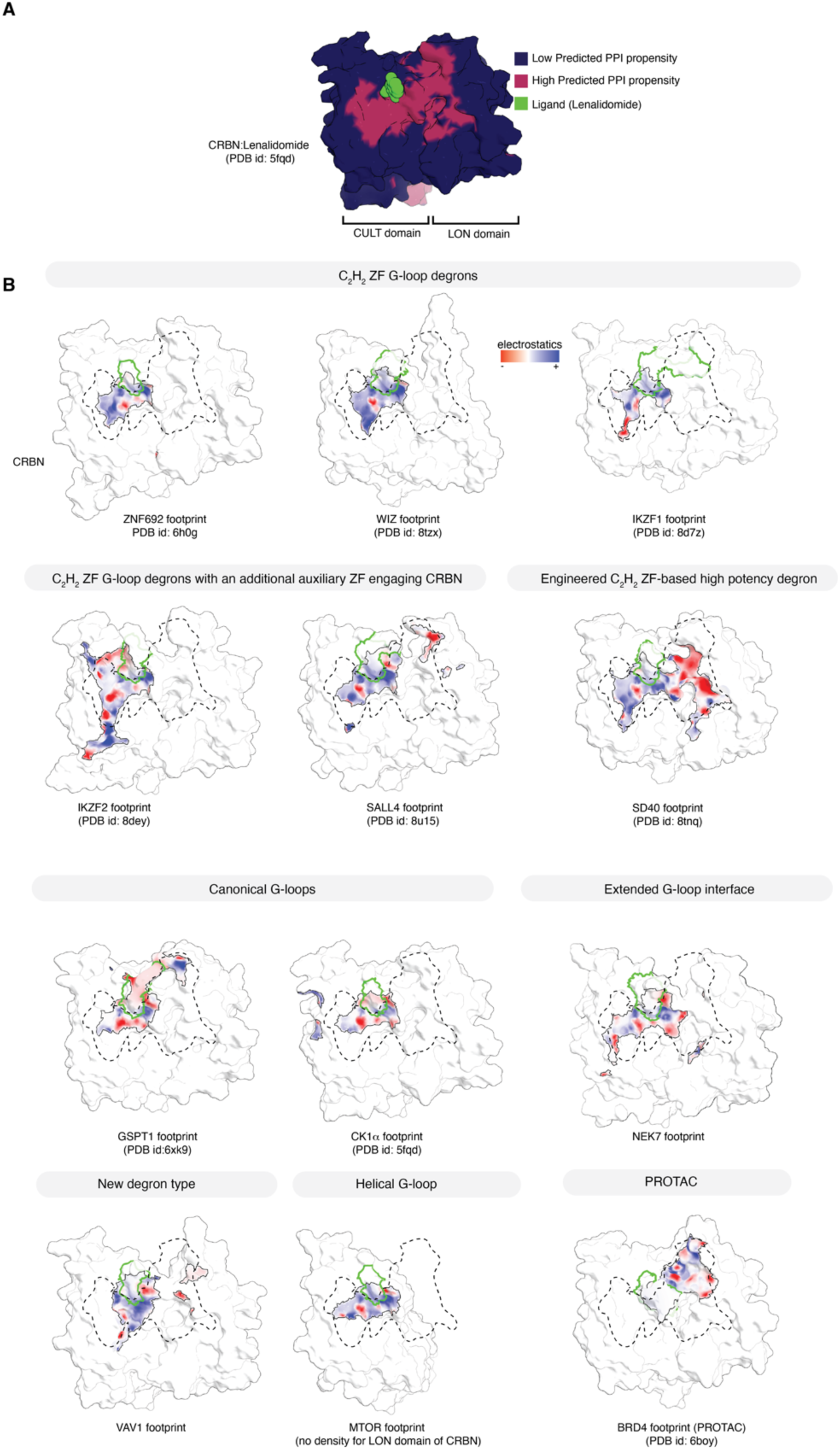
CRBN predicted PPI hotspot region and footprints of known neosubstrates. (A) Predicted PPI propensity at the surface of CRBN, with the molecular glue degrader shown in green as a reference. (B) Footprints of publicly available CRBN neosubstrates mapped on the CRBN:compound neosurface. The category of each neosubstrate is marked in the top bar. On each panel, the region that becomes buried upon engagement of the neosubstrate is colored according to the Poisson:Boltzmann electrostatics with a black line contour, while regions that are not buried upon neosubstrate engagement are shown in white. The surface region of the compound is marked by a green contour. A dotted contour line marks the predicted region of high PPI propensity on CRBN.

**Table S1.** Data collection and refinement statistics.

|  | DDB1/CRBN<sup>40-442</sup>-<br>Cpd. 6-VAV1(SH3c) | DDB1<sup>ΔB</sup>/CRBN<sup>40-442</sup>-Cpd. 1-mTOR-FRB | DDB1<sup>ΔB</sup>/CRBN<sup>40-442</sup>-Cpd. 5-NEK7 |
| --- | --- | --- | --- |
| **Data collection** |  |  |  |
| Space group | *P*6<sub>1</sub>22 | *P*2<sub>1</sub>2<sub>1</sub>2 | *P*2<sub>1</sub>22<sub>1</sub> |
| Cell dimensions |  |  |  |
| *a*, *b*, *c* (Å) | 143.60, 143.60,<br>367.64 | 183.18, 81.21,<br>99.02 | 118.58, 121.86,<br>144.74 |
| $\alpha$, $\beta$, $\gamma$ (°) | 90, 90, 120 | 90, 90, 90 | 90, 90, 90 |
| Resolution (Å)* | 47.4–3.40 (3.49–<br>3.40) | 74.2–2.95 (3.03–<br>2.95) | 85.1–3.25<br>(3.36–3.25) |
| *R*<sub>merge</sub> | 0.336 (>1.00) | 0.153 (>1.00) | 0.236 (>1.00) |
| *I*/σ*I* | 12.3 (1.1) | 10.9 (1.1) | 7.3 (1.2) |
| Completeness (%) | 98.2 (74.5) | 94.8 (60.2) | 74.3 (38.5) |
| Completeness [ellipsoidal] (%) | 98.2 (74.5) | 96.5 (72.4) | 94.3 (88.5) |
| Multiplicity | 38.2 (41.4) | 11.0 (6.8) | 13.1(13.0) |
| CC<sub>1/2</sub> | 0.998 (0.404) | 0.997 (0.438) | 0.995 (0.612) |
| **Refinement** |  |  |  |
| Resolution (Å) | 47.4–3.40 | 46.0–2.95 | 85.2–3.25 |
| No. reflections | 31052 | 29669 | 24342 |
| *R*<sub>work</sub> / *R*<sub>free</sub> | 0.213 / 0.271 | 0.240 / 0.261 | 0.212 / 0.259 |
| No. atoms (non-hydrogen) |  |  |  |
| Protein | 12071 | 9994 | 11511 |
| Ligand/ion | 23 | 21 | 69 |
| *B*-factors (Å<sup>2</sup>) |  |  |  |
| Protein | 131.7 | 80.8 | 94.5 |
| Compound | 137.8 | 80.0 | 111.8 |
| R.m.s. deviations |  |  |  |
| Bond lengths (Å) | 0.119 | 0.011 | 0.011 |
| Bond angles (°) | 1.33 | 1.31 | 1.31 |

\*Values in parentheses are for highest-resolution shell.

**Table S2.**
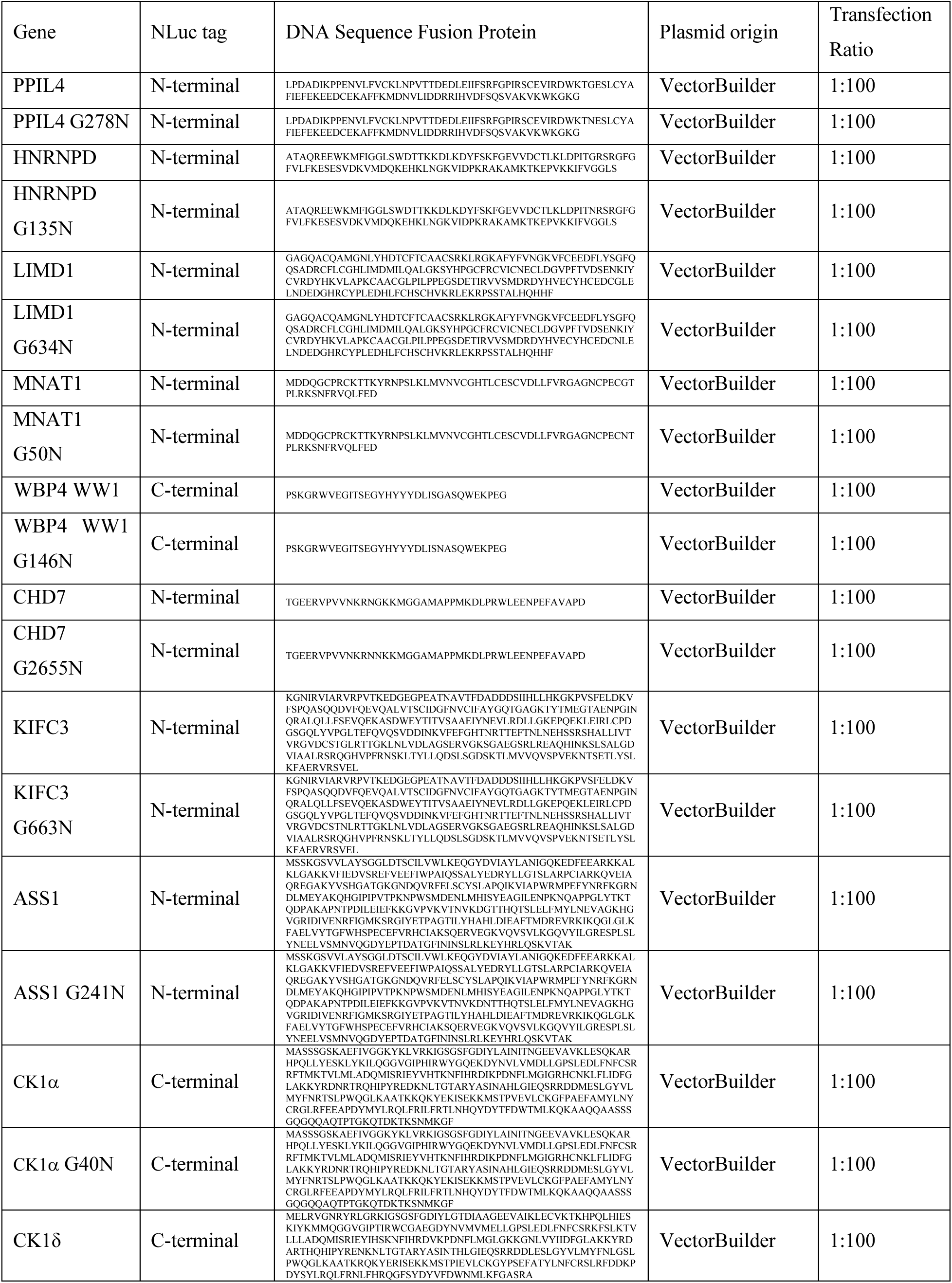
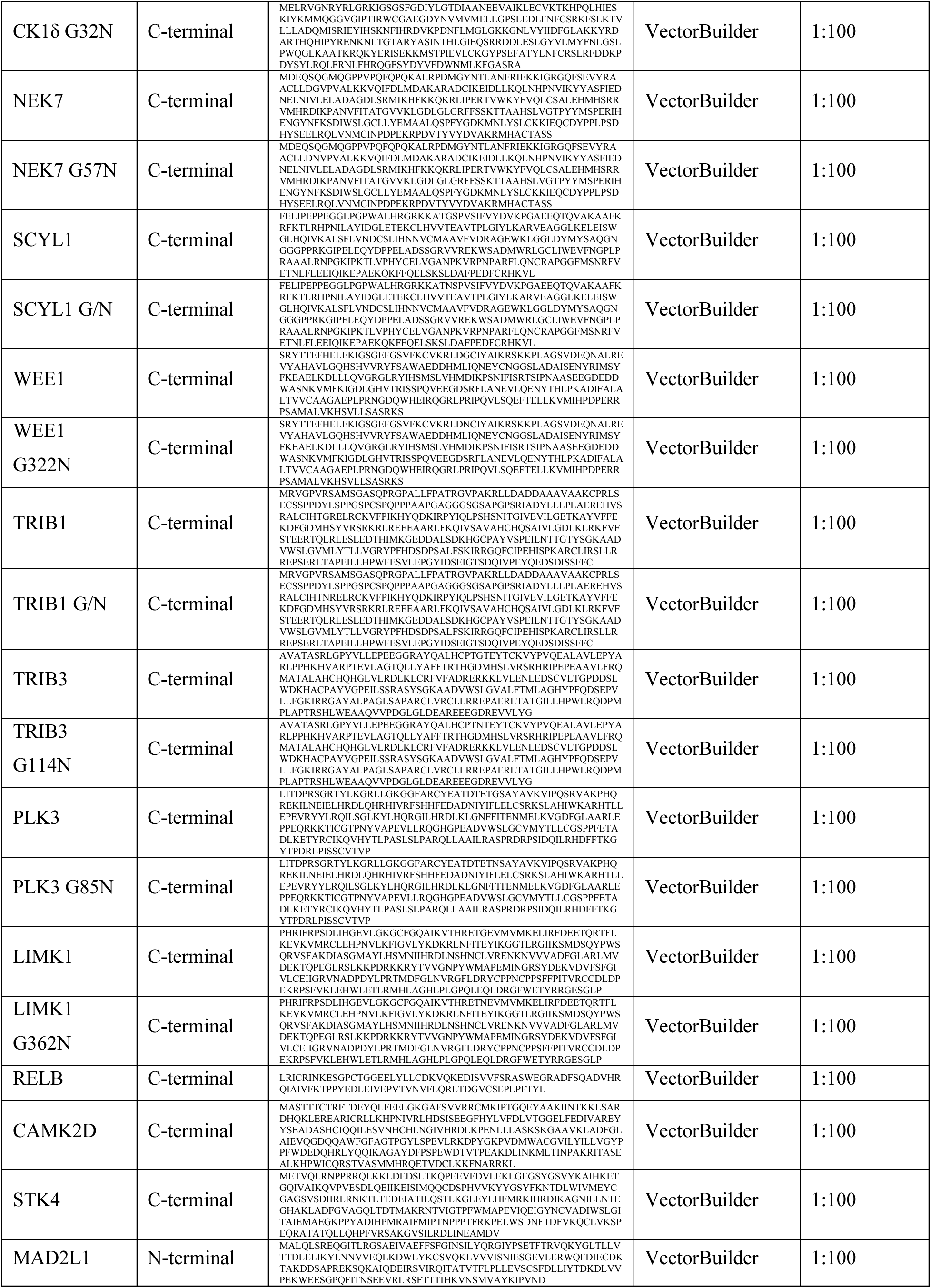
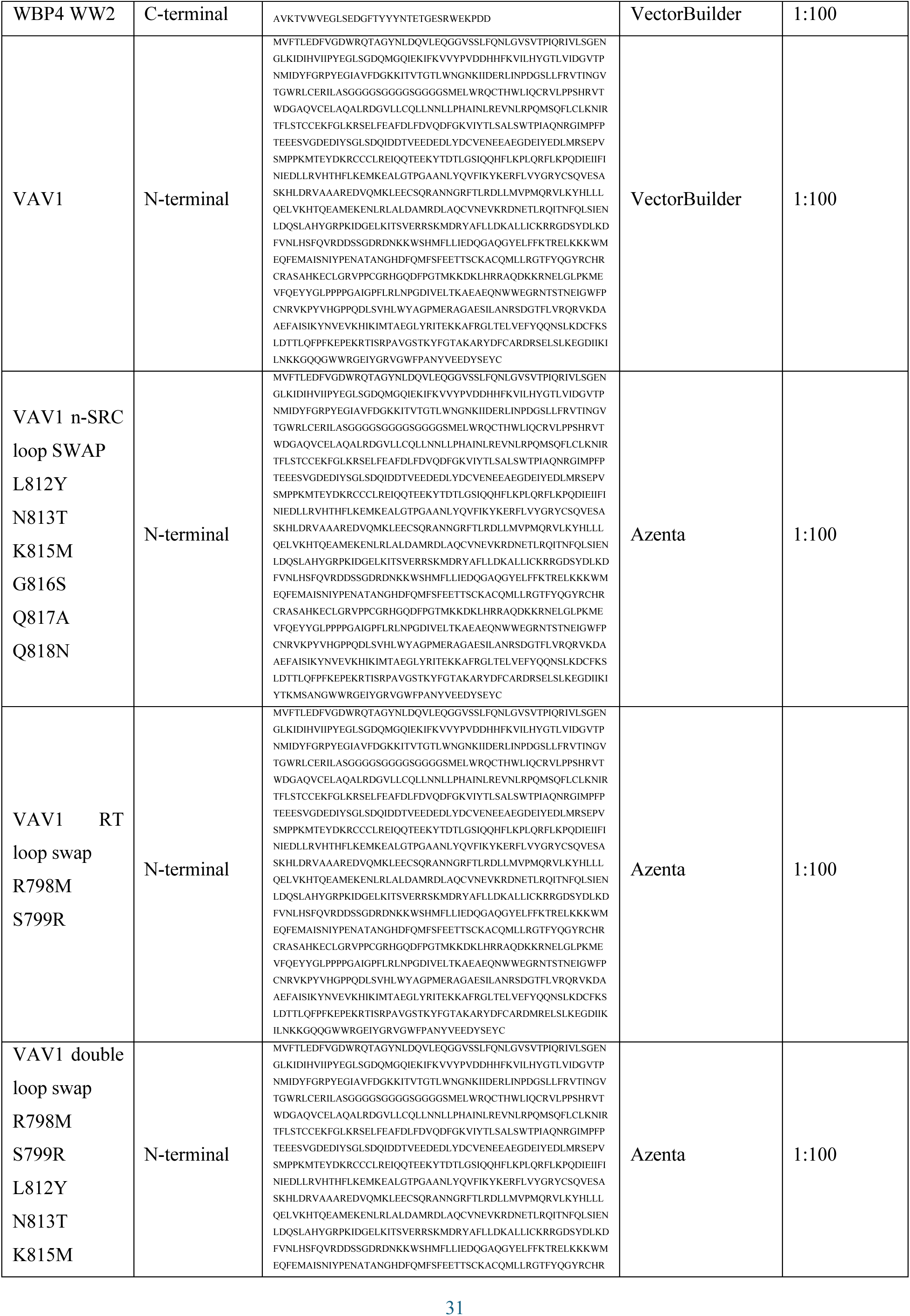
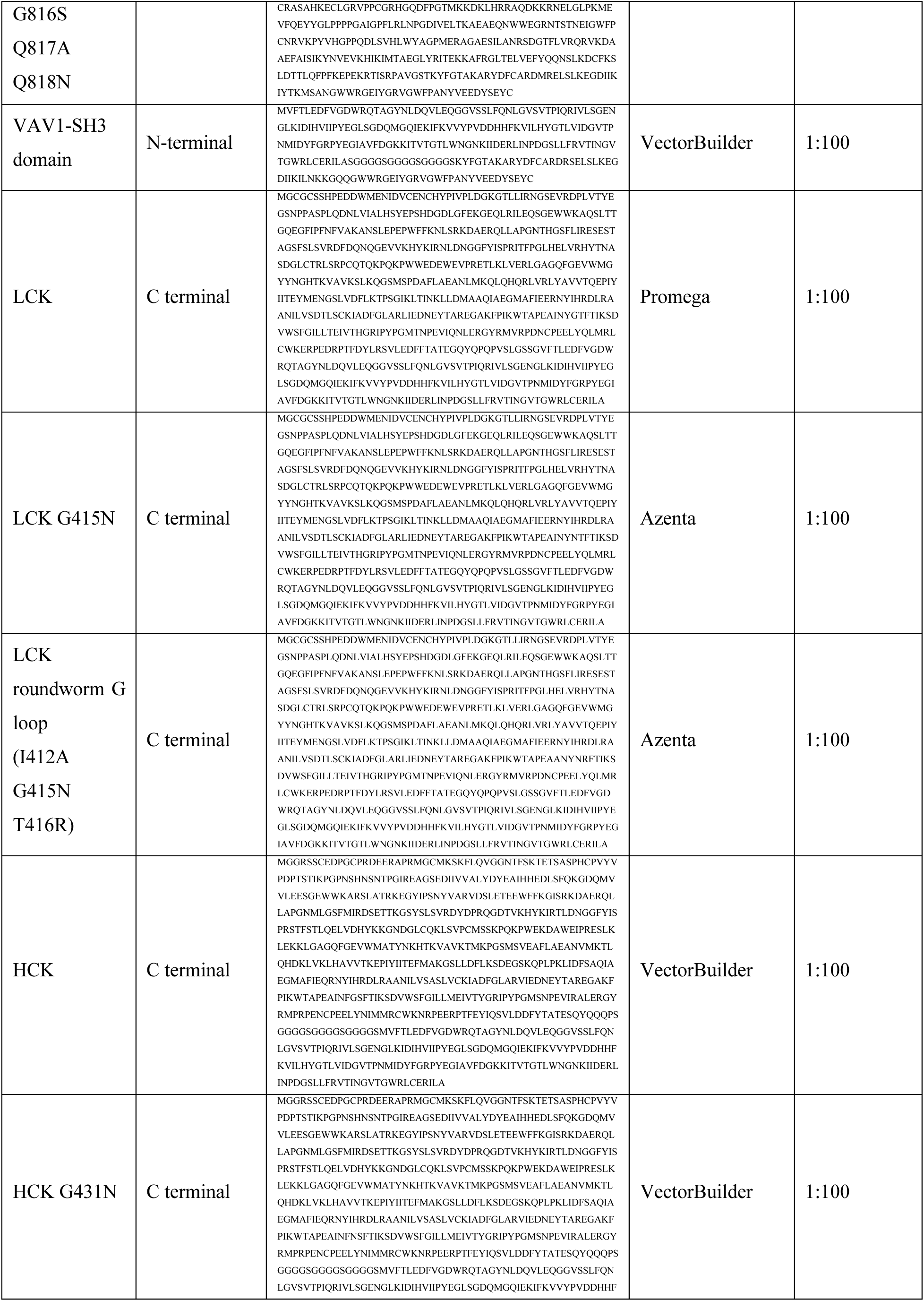
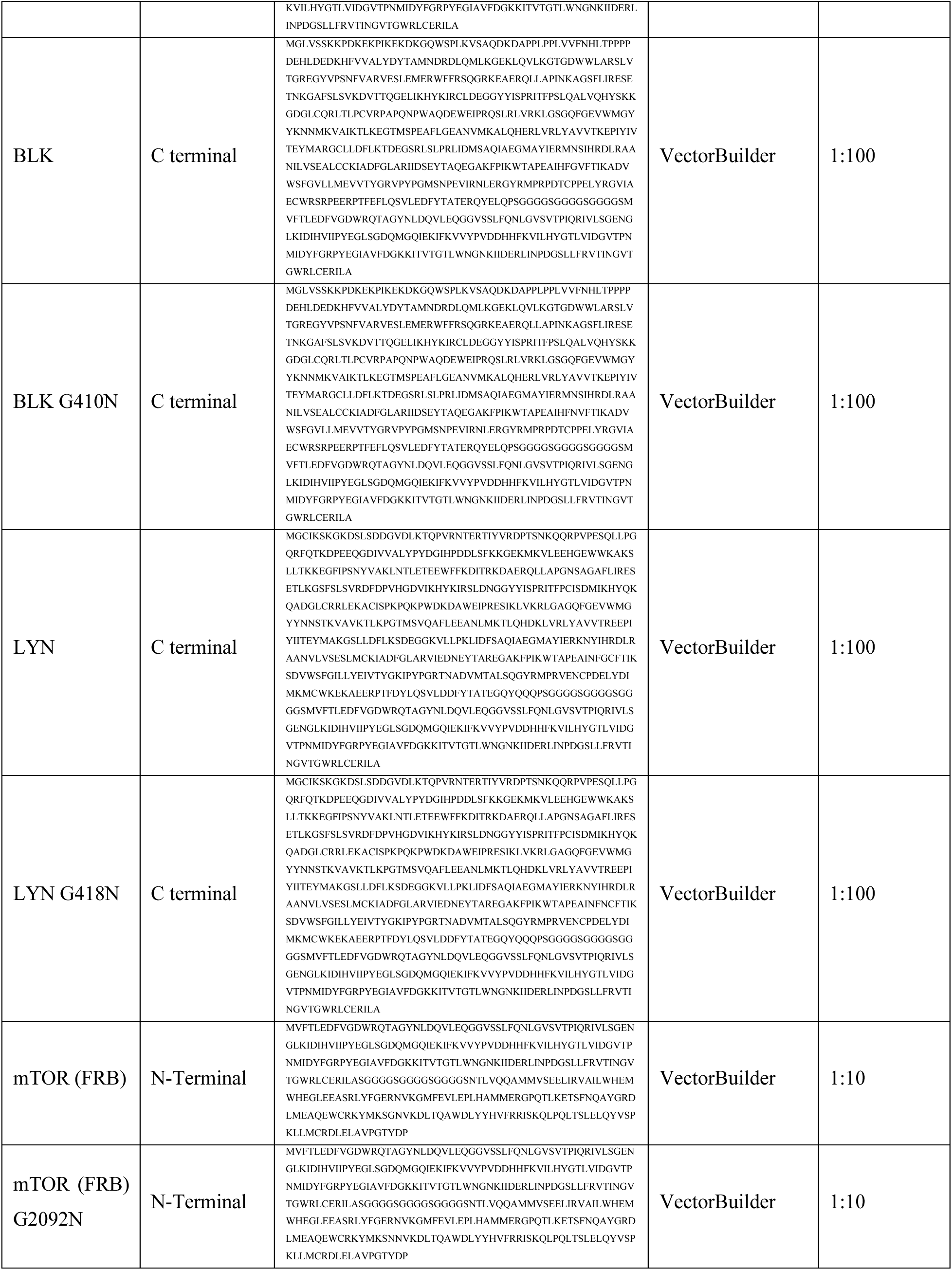
NanoLuc fusion vector constructs.

**Table S3.** Biochemical protein constructs.

| **Protein** | **Part #** | **Supplier** | **Affinity Tag** | **Boundaries** |
| --- | --- | --- | --- | --- |
| WEE1 | W01-11G-10 | SignalChem Biotech | N-terminal GST | 1-646 |
| SCYL1 | S04-35G-50 | SignalChem Biotech | N-terminal GST | 1-556 |
| TRIB1 | T30-30G-50 | SignalChem Biotech | N-terminal GST | 1-372 |
| PAK1 | P02B-10BG-10 | SignalChem Biotech | N-terminal GST | 1-545 |
| TRIB3 | T32-30G-20 | SignalChem Biotech | N-terminal GST | 1-358 |
| PLK3 | P43-13G-10 | SignalChem Biotech | N-terminal GST | 57-340 |
| CK1δ | C65-10G-10 | SignalChem Biotech | N-terminal GST | 1-415 |
| LIMK1 | L04-11G-10 | SignalChem Biotech | N-terminal GST | 285-638 |
| LCK | L03-10G | SignalChem Biotech | N-terminal GST | 1-509 |
| HCK | H02-11G | SignalChem Biotech | N-terminal GST | 230-497 |
| BLK | B02-10G | SignalChem Biotech | N-terminal GST | 1-505 |
| LYN | L13-18G | SignalChem Biotech | N-terminal GST | 1-512 |
| NEK7 | N09-10G | SignalChem Biotech | N-terminal GST | 1-302 |

## Materials and Methods

### Mining of the human proteome for *β-hairpin* and helical G-loops

A bioinformatic pipeline was employed to systematically mine human proteome structural datasets for β-hairpin or helical G-loop motifs (referred to as G-loop here) associated with CRBN-binding, leveraging both experimentally solved protein structures from the Protein Data Bank (PDB) and AlphaFold2 (AF2) models. The CK1α G-loop (PDB ID: 5FQD) was used as a template motif, defined by a conserved glycine residue (G40) positioned within an eight-residue β-hairpin α-turn around G40, spanning I35 to E42 or a minimal five-residue motif spanning I37 to I41.

#### 1. Structural Data Preparation

All protein chains corresponding to human proteins in the Protein Data Bank (PDB) were extracted from the deposited structures, and all non-protein atoms were removed using the Biopython PDB manipulation library (*48*). As the human proteome is incompletely covered in the PDB, annotated structural domains in Alphafold2 (AF2) models for human proteins were included in the dataset. Specifically, PFAM (*49*), Prosite (*50*) or SMART (*51*) domain ranges for all human genes standard Uniprot genes were extracted from the InterPro database (*52*). The start and end ranges for these domain definitions was then used to extract substructures from the AF2 v4 database (*53*), and each substructure was saved as a separate file.

#### 2. G-loop Identification and Superposition

Motifs resembling the CK1α G-loop were identified using a sliding window approach that traversed each protein’s sequence and checked for the presence of glycine at a specific position. Continuous peptide segments of length equal to the template G-loop were extracted. Motifs containing a glycine at the central position were subjected to structural alignment with the template G-loop using the Superimposer module from Biopython library (*48*). RMSD values between aligned motifs were calculated and at this filtering stage motifs with an RMSD below 1.0 Å were selected for further processing, independent of the template G-loop length.

#### 3. Clash Filtering with CRBN

Once the protein chain structure containing a 5- or 8- residue G-loop was aligned to the template CK1α G-loop, we assessed potential steric clashes between the aligned structure and the closed CRBN conformation found in the CK1α structure (PDB ID: 5FQD). Clashes were defined as instances where the distance between backbone atoms of the aligned motif and any heavy atom of CRBN was less than 2.0 Å. Motifs with fewer than 12 total clashes and fewer than 1 clash involving the Cα atoms of the matched domain were retained. The secondary structure of each motif was then assigned using the DSSP program (*54, 55*) for further analysis.

#### 4. Postprocessing of matched motifs

After processing, a domain annotation was assigned to each structural chain where the G-loop was contained. Specifically, PFAM (*49*), Prosite (*50*) or SMART (*51*) domain ranges for all human genes standard Uniprot genes were extracted from the interprosite database and assigned to the matched G-loop. The PANTHER classification system (*56*) was used to classify the gene of each G-loop to a class. At this point, G-loops were further filtered according to their RMSD with respect to the template. In the 8-residue template mining pipeline, G-loops with an RMSD > 0.75 Å with respect to the template CK1α G-loop were discarded from consideration at this point, while in the minimal 5-residue pipeline, G-loops with an RMSD > 0.5 Å were discarded. As many of the identified G-loops appear in domains that are often found only in extracellular proteins, G-loops in these domains were discarded from consideration.

The dataset of predicted G-loops in the human protein with the 8-residue window is available in a csv format in **Data S1**, while the dataset with the minimal 5-residue window is available in **Data S2**.

### Surface computation and visualization

The MaSIF program(*57, 58*) was used to compute all surfaces used in the paper, including electrostatics. MaSIF uses the PyMesh library (*59*) to process surface meshes, and Adaptive Poisson-Boltzmann Solver (APBS) (*60*) to compute electrostatics.

#### 1. Definition of buried surface areas

We used the definition of buried surfaces (i.e. the interface of a PPI) identical to that presented in (*57, 58*). PyMesh was applied to regularize the meshes at 1.0 Å resolution. The buried surface of a subunit in a complex structure was calculated by comparing surface vertices of the subunit’s surface mesh and the complex’s surface mesh. Vertices in the subunit’s mesh were labeled as ‘interface’ meshes if they were more than 2.0 Å away from any complex surface vertex.

#### 2. Prediction of CRBN PPI propensity

The MaSIF-site program (*57*) was used to compute a per-vertex propensity of the CRBN surface (PDB id: 5fqd) to become buried in a PPI, as visualized in Fig. S10A. MaSIF-site uses as input the per-vertex chemical and geometrical features projected on the surface of a protein, and uses them to compute a score for each vertex shown to correspond to its propensity become buried in a protein-protein interaction (*57*). MaSIF-site is built on a geometric deep learning architecture trained to predict the buried surface from a training set of thousands of proteins. The training set did not contain the CRBN protein. For visualization purposes, the propensity of each vertex, computed as a real value between 0 and 1, was rounded to the nearest integer.

#### 3. Surface matching of the VAV1 similarity

The MaSIF-seed program (*57, 58*) is a surface matching pipeline that uses geometric deep learning to encode surface patches as fingerprints. MaSIF-seed was designed and tested to match surfaces that are complementarity to each other based on their fingerprint, followed by an alignment step. MaSIF-seed was modified to match patches by similarity (or *mimicry*) instead of complementarity.

#### 4. Overall method description of MaSIF-seed

MaSIF-seed method is a geometric deep learning approach designed to identify complementary structural motifs that can mediate protein-protein interactions. It works by analyzing protein molecular surfaces, which are divided into overlapping radial patches, to capture both chemical and geometric features relevant to binding. Every vertex on the surface becomes the center of its own radial patch of radius 12 Å. A neural network is then trained to generate “fingerprints” for these overlapping surface patches that predict their binding propensity. MaSIF-seed uses this fingerprint matching process to find complementary surface motifs from a large database of potential binding partners. The MaSIF-seed method involves an important alignment step after the initial identification of complementary surface motifs. Once the surface patches from the target and potential binding seeds are matched based on their fingerprints, these patches are aligned geometrically to evaluate how well they fit together. A second neural network, called the interface post alignment (IPA) score network is used to score the aligned surface patches specifically for complementarity.

#### 5. Definition of the CK1α and GSPT1 degrons as a surface patch

The molecular surfaces of CK1α (PDB id: 5fqd) and GSPT1 (PDB id: 6xk9) from their co-crystal structures in complex with CRBN were used as input to the algorithm. Their buried surface in the ternary complex with CRBN/MGD was computed as described above. The degron patch was defined as the patch of radius 12 Å that captures the largest area of the degron region.

#### 6. Modification of MaSIF-seed for surface similarity

MaSIF-seed was modified to match and align similar surface patches instead of complementary patches. MaSIF-seed by default inverts the input features and normals of the target patch to align for complementarity, but this is no longer required for similarity matching. To align patches by similarity, these features were no longer inverted. MaSIF-search fingerprints were computed for all the patches decomposed from the molecular surfaces of VAV1 domains (Supp. Fig. 9A). In a first stage patches with fingerprints with an Euclidean greater than 2.5 were discarded. Patches that passed this filter were selected for a second stage alignment step. Since every vertex in the surface is the center of a patch, each vertex on the surface has a computed fingerprint. In the second stage alignment step, the RANSAC algorithm is used to perform an alignment between the patches. In each iteration of RANSAC three vertices in the matched patch are randomly selected, and the most similar fingerprint in the query patch (GSPT1 or CK1α) are identified. The three matches are used to perform an alignment and the number of inlier vertices after the alignment is used to initially score the alignment. The alignment with the highest number of inliers is accepted. Finally, a scoring of similarity is performed using the scored fingerprints. Specifically, after an alignment all vertices of the aligned patch within a 3D distance of 2.0 Å from a vertex in the query patch are selected. The fingerprint distance *d* is computed between the selected vertices and each of the corresponding nearest neighbor’s fingerprint in the query patch. The score for the alignment is computed as 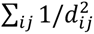 such that *i* is the vertex in the aligned patch and *j* is the vertex in the query patch and *d* is the fingerprint Euclidean distance. The interface post-alignment (IPA) scoring neural network (*58*) was set to a minimum value (0.20) and its score ignored as it is not applicable to this problem. MaSIF-seed contains a steric, atomic based post-processing filter which is used by default to filter matched results for steric compatibility with the complementary target. Since similarity was performed here, the input was modified so that the steric filter comparison is performed against closed CRBN in the respective structure (PDB id: 5fqd or 6xk9), removing any solution where no steric clashes involving Cα atoms were allowed, and up to 10 clashes involving other heavy atoms were allowed.

#### 7. Computation of surface similarity

To visually display the surface similarity of GSPT1 and VAV1 after the surface-based alignment and scoring, a per-surface-vertex surface similarity was computed on the surfaces after alignment by the algorithm. The surface similarity score was computed in the same way that per-vertex shape complementarity has been previously defined (*61*). Let *A* and *B* be two proteins aligned under our surface alignment algorithm. Let *P_A_* be the molecular surface of protein *A*, and *x_A_* be a vertex on the surface of *A* with normal *n_A_*. Let *P_B_* be the molecular surface of protein *B*, and 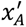 be the point on the surface of *B* closest to *x_A_* with normal 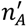. Then, the surface similarity for vertex *x_A_* is defined

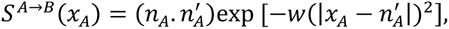

where *w*=0.25.

### Mass spectrometry-based methods

CAL-51 cells were genetically modified by transduction with lentiviral particles derived from the construct cLV-CMV-TurboID-hCRBN-IRES-PuroR, which contains a puromycin resistant gene to select cells expressing the TurboID CRBN fusion. The multiplicity of infection (MOI) defined as the ratio of cells surviving selection over cell number in the no virus/no puromycin control was 0.5. CAL-51 TurboID CRBN cells were cultured in DMEM (Cat.No. 11965-092, Gibco) supplemented with 10% FBS (Cat# P30-1909, PAN Biotech) and 2 µg/mL puromycin (Gibco, #A11138-03).

#### 1. Cell Treatment for TurboID or Global Proteomics Experiments

For TurboID experiments, 6-well plates of sub confluent 70-90% CAL51 cells were incubated for 6h at 37°C in fresh medium DMEM high Glucose (Gibco, 11965) + 10% FBS (PAN Biotech, P30-109) with 50 µM D-biotin (Thermo Scientific, B20656), 2 µM of Bortezomib (Selleckchem, S1013) and 10 µM compound or DMSO (Fisher Bioreagents, BP231-100). For Global Proteomics profiling experiments, 12-well plates of sub confluent 70-90% CAL51 cells were incubated for 24h at 37°C in fresh medium DMEM high Glucose (Gibco, 11965) + 10% FBS (PAN Biotech, P30-109) with 10 µM compound or DMSO (Fisher Bioreagents, BP231-100). Cells were washed once with PBS and collected with 0.2mL of TrypLE (Gibco, # A12177) in 1.5 mL Eppendorf tube lowbind. Cells were pelleted by centrifugation at 500xg for 4min at 4°C, supernatants were removed, and cells were washed with 1mL of PBS. After centrifugation, dried pellets were frozen to –80°C.

#### 2. TurboID Sample Preparation

Cell pellets were lysed in 90uL of cold RIPA lysis buffer (25 mM Tris-HCl pH 7.6, 150 mM NaCl, 1% NP-40, 1% sodium deoxycholate, 0.1% SDS; Thermo Scientific, 89901), protease inhibitors (Sigma, I3786, P5726, P0044), 2 µM Bortezomib (Selleckchem, S1013), then sonicated at 4°C using the QSonica Q800R3 for 3min using 80% amplitude for 5s pulse, and 2s pause to disrupt visible aggregates. Cell lysates were transferred into a 96-well plate and incubated with 30 µL of pre-washed streptavidin beads (in RIPA-buffer) for 4h at 4°C. The BRAVO automation system from Agilent equipped with a magnetic plate holder was used to wash the beads 2 times with 140 µL RIPA buffer, 2 times with 140 µL 2M urea (Sigma, U1250) 10mM Tris-HCl pH 8.0 (Alfa Aesar, J22638), and 2 times with 140 µL cold PBS. The washed beads were then incubated with 25 µL PreOmics iST-NHS lysis buffer (PreOmics, Cat. No. iST-NHS96x P000030) and processed using the PreOmics kit following their recommended protocol with minor modifications. In brief, the proteins were reduced, alkylated and digested for 3 hr at 37 °C. The peptides were then labelled with tandem mass tag (TMT) reagent (1:4; peptide:TMT label) (Thermo Fisher Scientific). After quenching, the peptides from the 16 conditions were combined to a 1:1 ratio and purified. Mixed and labeled peptides were subjected to high pH reversed-phase fractionation with Pierce^TM^ High pH Reversed-Phase Peptide Fractionation Kit (Thermo Fischer, #84868), according to their recommended protocol. After drying in a speed vac, each 7 fractions of labelled peptides were loaded onto an Aurora column from Ionopticks (75 µm ID, 1.6 µm particles, 25 cm in length) in an EASY-nLC 1200 system. The peptides were separated using a 168 min gradient from 4 % to 30 % buffer B (80% acetonitrile in 0.1 % formic acid) equilibrated with buffer A (0.1 % formic acid) at a flow rate of 400 nl/min. Eluted TMT peptides were analyzed on an Orbitrap Eclipse mass spectrometer (Thermo Fisher Scientific).

#### 3. Global Proteomics Profiling Sample Preparation

Cell pellets were lysed using 75 µL PreOmics iST-NHS lysis buffer. A tryptophan assay was used to determine lysate concentration. Plates were read with a Tecan Infinite 200 PRO and lysates were then normalized to 100 μg in 50 μL. The normalized lysates were then processed using the recommended PreOmics sample preparation protocol with minor modifications. In brief, proteins were reduced, alkylated, and digested for 3 hrs at 37 °C. The peptides were then labelled with tandem mass tag (TMT) reagent (1:4; peptide:TMT label) for 1 hr at 22 °C. Labeled peptides were then quenched with 5% hydroxylamine solution diluted in LC/MS-grade water. The peptides from the 18 conditions were then combined in a 1:1 ratio and purified with PreOmics desalting cartridges. Mixed and labeled peptides were subjected to high-pH reversed-phase HPLC fractionation on an Agilent X-bridge C18 column (3.5 µm particles, 2.1 mm I.D, and 15 cm in length). Using an Agilent 1200 LC system, a 61 min linear gradient from 0% to 45% acetonitrile in 10 mM ammonium formate, pH 10, separated the peptide mixture into a total of 60 fractions, which were then consolidated into 24 fractions containing an assortment of TMT-labeled fragments in each. Peptides concentrations were estimated by UV at 14.58 ug/uL per fraction. After drying in a SpeedVac integrated vacuum concentrator, all 24 fractions from the plate were resuspended in the same volume of 0.1% formic acid calculated by utilizing the chromatogram generated by the HPLC. Each of the fraction was loaded onto a 25 cm Aurora column (75 µm I.D., 1.6 µm particles) using a Vanquish Neo HPLC system. The peptides were separated using a 168 min gradient from 4% to 30% buffer B (80% acetonitrile in 0.1% formic acid) equilibrated with buffer A (0.1% formic acid) at a flow rate of 400 nL/min. Eluted TMT peptides were analyzed using an Orbitrap Eclipse mass spectrometer.

#### 4. Data Acquisition and Analysis

MS1 scans were acquired at resolution 120,000 with 400-1400 mass over charge (m/z) scan range, Automatic Gain Control (AGC) target 4 x 10^5^, maximum injection time 50 ms. Then, MS2 precursors were isolated using the quadrupole (0.7 m/z window) with AGC 1 x 10^4^ and maximum injection time 50 ms. Precursors were fragmented by collision-induced dissociation (CID) at a normalized collision energy (NCE) of 35% and analyzed in the ion trap. Following MS2, synchronous precursor selection (SPS) MS3 scans were collected by using high energy collision-induced dissociation (HCD) and fragments were analyzed using the Orbitrap (NCE 55 %, AGC target 1 x 10^5^, maximum injection time 120 ms, resolution 60,000). Protein identification and quantification were performed using Proteome Discoverer 2.5.0.400 with the SEQUEST algorithm and UniProt human database (2021-01-29, 20614 protein sequences). Mass tolerance was set at 10 ppm for precursors and at 0.6 Da for fragments. A maximum of 3 missed cleavages were allowed. Methionine oxidation was set as dynamic modification, while TMT tags on peptide N termini/lysine residues and cysteine alkylation (+113.084) were set as static modifications. Adjustment of reporter ion intensities for isotopic impurities according to the manufacturer’s instructions was performed in Proteome Discoverer. The list of identified peptide spectrum matches (PSMs) was filtered to respect a 1% false discovery rate. Furthermore, PSMs with an average TMT reporter ion signal-to-noise value lower than 10, a precursor isolation interference level value higher than 50%, and with any empty TMT channels were excluded. Subsequently, protein identifications were inferred from unique peptides, i.e. peptides matching multiple protein entries were excluded. Protein relative quantification was performed using an in-house developed software in Java and R. This analysis included multiple steps; filtering PSMs (as described above), global data normalization by equalizing the total reporter ion intensity across all channels, summation of reporter ion intensities per protein and channel, calculation of protein abundance L2FC and testing for differential abundance using moderated t-statistics (R LIMMA (Phipson et al., 2016)) where the resulting p-values reflect the probability of detecting a given L2FC across sample conditions by chance alone. An arbitrary L2FC cut-off of −1 and +1 was applied to the dataset corresponding to a minimal −50% depletion (x0.5) or +100% enrichment (x2) respectively. The p-value cut-off for statistical significance was set at the standard 10^-2^ value.

### Cellular NanoBRET

The mammalian gene expression vector for transient transfection of HaloTag®-CRBN was purchased from Promega. NanoLuciferase (NanoLuc)-Fusion vectors were either purchased or obtained by mutagenesis from VectorBuilder, Azenta, or Promega. The NanoLuc-fusion vectors were designed consisting of either C terminal or N terminal NanoLuc fused to a human target gene trough a 3xGGGGS or VSLGSSG linker. HEK293 cells were cultured in Dulbecco’s modified Eagle’s medium (DMEM) (Gibco) +10% fetal bovine serum (FBS) (Gibco). Medium was removed from flasks by aspiration, cells were washed with phosphate buffered saline (PBS), and trypsin-EDTA was added to dissociate cells. Culture medium was added to the cells to neutralize trypsin, then cells were transferred in a centrifuge tube and centrifuged at 300×g for 3 minutes. Media was aspirated from the cell pellet and cells were resuspended in culture medium and passed through a 40-μm cell strainer. Cells were counted using a Cellometer® cell counter and then diluted to 4×10^5^ cells/mL in full media. Cells were seeded in 6 well plates, 2 mL/well for a total 8×10^5^ cells/well. Cells were incubated overnight at 37°C, 5% CO2. On day two, cells were transfected with a Nanoluc (Donor): HaloTag (Acceptor) DNA co-transfection mix using Fugene HD (Promega). Donor: acceptor vector ratio of 1:1, 1:10 or 1:100 was selected depending on target (see supplemental materials) HaloTag-CRBN and Target-NanoLuc fusion vectors were added to 100 μL phenol red-free OptiMEM (Gibco). Finally, 6 μL FuGENE HD was added to the transfection mix, and the mix was gently agitated. After incubation for 10-15 minutes at room temperature, each transfection mix was added dropwise to one well of the overnight-cultured cells. Multiple wells were transfected for each target to perform the experiment. Cells were incubated overnight at 37°C, 5% CO2. After 18-20 hours, media was aspirated, cells were washed with PBS, and then trypsin was added. Dissociated cells were collected using phenol red-free OptiMEM +4% FBS, placed in a centrifuge tube, and centrifuged at 300×g for 3 minutes. Media was then aspirated from the pellet, and the pellet was resuspended in phenol red-free OptiMEM + 4% FBS. Cells were counted as previously described and diluted in phenol red-free OptiMEM +4% FBS to 1.5×10^5^ cells/mL. HaloTag 618 Ligand (Promega) was added to the cells. Cells treated with DMSO and without 618 ligand were used to calculate background. Cells were seeded in a white, solid bottom 384 well plate (Corning# 3570), 25 μL/well. Plates were then incubated overnight at 37°C and 5% CO2. On day four, compounds were added to the plates in 10-point, 1:3 dilution series starting at 10 μM using an Echo Acoustic Liquid Dispenser. DMSO was transferred to all the control wells. After treatment, plates were returned to the incubator for 3 hours. After three hours, NanoBRET Nano-Glo Substrate (Promega) was diluted in Phenol red-free OptiMEM +4% FBS and added to the plates using a multidrop Combi. Dual luminescence measurements were then taken on a PHERAstar® FSX using the LUMI 610 LP 450-80 H optic module. The ratio between signal recorded in the acceptor channel and the donor channel was used to determine the Bioluminescence Resonance Energy Transfer (BRET) at a well-to-well level. Average signal from wells containing no 618 ligand was used to subtract background and curves were normalized to the average DMSO treated wells containing 618 ligand. For Figures 1, 2, and 3, NanoBRET curves were further normalized, such that the values were rescaled to a range between 0 and 1 by subtracting the minimum and dividing by the range of the data (maximum – minimum). Correspondingly, the standard deviations were normalized by dividing by the same range factor.

### TR-FRET assay

Kinase proteins used in TR-FRET experiments (WEE1, SCYL1, TRIB1, PAK1, TRIB3, PLK3, CK1δ, LIMK1, LCK, HCK, BLK, LYN, and NEK7), were sourced commercially from SignalChem Biotech (Table S3), others were produced as described in the protein expression and purification sections. A one-pot detection solution of 6His-tagged CRBN/DDB1 complex (12.5 nM), GST-tagged neosubstrate (25 nM), anti-GST-d2 (1X), and anti-6xHis Tb Cryptate Gold (1X) was prepared in 20 mM HEPES, 20 mM NaCl, 0.2 mM TCEP, 0.2 mM EDTA, and 0.005% Surfactant P20 and was dispensed to each assay plate. Biochemical assays were conducted in 384-well white Proxi plates in 20 µL total volume. MGD was stored in dry, ambient temperatures at 10 mM. A 10-point, 1:3 dilution series was prepared from 10 mM stock concentrations in Echo-compatible low dead volume plates. A volume of 20 nL of the MGD dilution series was dispensed into assays wells using an Echo 650 (Beckman, USA). A volume of 20 nL of dimethyl sulfoxide was transferred into the neutral-control wells. The assay was then allowed to incubate for 45 minutes at ambient temperature after transferring compound. Plate measurements were taken on a PHERAstar FSX using the HTRF Red emission filter (example: 337 nm, em1: 620 nm, em2: 665 nm; Flashes: 27, Integration time: 60-400 us, Z-height: 10 mm, Ratio-multiplier: 10,000).

### Protein expression and purification

#### 1. Expression and Purification of CRBN-B02/DDB1-ΔB-B06

The MRT-CRBN-B02 DDB1-B06 complex was expressed using the Bac-to-Bac Baculovirus Expression System in *High Five* (HF) cells. The N-terminally truncated MRT-CRBN-B02(Δ1-40) construct (residues 41–442) included a StrepII tag followed by a TEV cleavage site. MRT-DDB1-B06 was expressed with deletion of the BPB of DDB1, with N-terminal residues 1-395 and the C-terminal residues 706-1140 from DDB1. The expression was performed in a 5 L culture with a multiplicity of infection (MOI) of 10 and supplemented with 20 µM ZnCl₂. The cell pellet was resuspended lysis buffer (50 mM Tris-HCl, pH 8.0, 150 mM NaCl, 1 mM EDTA, 10% glycerol, 1 mM PMSF, protease inhibitors, and Benzonase) and lysed by homogenization. Clarified lysate was incubated with 50 mL Strep-Tactin resin, washed, and eluted using 50 mM Tris-HCl, pH 8.0, 150 mM NaCl, 1 mM EDTA, 10% glycerol, and 50 mM D-biotin. The eluted protein was split, and a fraction was dialyzed in 5 L buffer (50 mM Tris-HCl, pH 8.0, 100 mM NaCl, 1 mM EDTA, 15% glycerol) at 4°C overnight with His-tagged TEV protease (1:30, w/w). Cleaved protein was purified using reverse Strep-Tactin affinity chromatography. The cleaved and uncleaved protein fractions were separately diluted to 80 mM NaCl and loaded onto a Q HP 5 mL column, then eluted with a gradient of 8%- 100% Buffer B (50 mM MES, pH 6.5, 1 M NaCl, 5% glycerol, 1 mM TCEP) over 30 column volumes. Fractions were pooled based on SDS-PAGE. The pooled fractions were concentrated and loaded onto a Superdex 200 16/600 column equilibrated with SEC buffer (10 mM HEPES, pH 7.4, 150 mM NaCl, 1 mM TCEP). Fractions containing the target complex were pooled, concentrated, flash-frozen in liquid nitrogen, and stored at −80°C.

#### 2. Expression and Purification of CRBN-B02/DDB1-B05

The coding sequences for MRT-CRBN-B11 (human CRBN residues 40-442 with an N-terminal StrepII-TEV tag) and MRT-DDB1-B05 (human DDB1 full-length residues 1-1140) were cloned into separate pFastBac1 vectors. Sf9 cells were infected with a multiplicity of infection (MOI) of 10 and incubated at 27°C for 48 hours. The harvested cell biomass was thawed on ice and was resuspended in in lysis buffer (100 mM Tris-HCl, pH 8.0, 150 mM NaCl, 1 mM EDTA, 5% glycerol, 1 mM PMSF, 1 tablet of cOmplete™ EDTA-free protease inhibitor per 200 mL, and 250 U/mL Benzonase®). Cells were lysed using a high-pressure homogenizer, initially at 200 bars for two cycles and then at 450 bars for four additional cycles. The lysates were clarified by centrifugation at 20,300 xg for 90 minutes at 4°C. The clarified lysates were incubated with Strep-Tactin beads pre-equilibrated with binding buffer (100 mM Tris-HCl, pH 8.0, 150 mM NaCl, 1 mM EDTA, 5% glycerol) overnight at 4°C. The resin was packed into a gravity flow column and washed with binding buffer. The bound proteins were eluted using elution buffer (100 mM Tris-HCl, pH 8.0, 150 mM NaCl, 1 mM EDTA, 5% glycerol, 50 mM D-biotin).TEV protease was added to the elute and the sample was dialyzed overnight at 4°C in dialysis buffer (100 mM Tris-HCl, pH 8.0, 150 mM NaCl, 1 mM EDTA, 15% glycerol). Following TEV cleavage, the protein sample was incubated with Strep-Tactin beads for 2 hours to remove the StrepII tag.The eluted proteins were diluted with binding buffer to reduce the NaCl concentration to 15 mM. The diluted sample was loaded onto a Q HP 5 mL column (Cytiva) equilibrated with Q-buffer A (50 mM MES, pH 6.5, 5% glycerol, 1 mM TCEP). The column was washed with 10 column volumes (CV) of 2% Q-buffer B (50 mM MES, pH 6.5, 1 M NaCl, 5% glycerol, 1 mM TCEP), followed by a 30 CV gradient from 2% to 100% of Q-buffer B to elute the proteins. Eluted fractions (6–15) were pooled and concentrated using an Amicon Ultra centrifugal filter (30 kDa MWCO) at 3000 xg at 4°C. Pooled fractions from the IEX step were concentrated to approximately 3.5 mL, centrifuged at 20,000 xg to remove any precipitates, and loaded onto a Superdex 200 16/600 PG column equilibrated with SEC buffer (10 mM HEPES, pH 7.4, 150 mM NaCl, 1 mM TCEP). The column was run at a flow rate of 1 mL/min, and fractions (9–21) were collected, pooled, and concentrated. The purified MRT-CRBN-B02/DDB1-B05 complex was aliquoted and snap-frozen in liquid nitrogen before storage at −80°C.

#### 3. Expression and Purification of CRBN-B11/DDB1-B05

The coding sequences for MRT-CRBN-B11 (human CRBN residues 40-442 with an N-terminal His6-TEV-SM tag) and MRT-DDB1-B05 (human DDB1 full-length residues 1-1140) were cloned into separate pFastBac1 vectors. Recombinant bacmids were generated using the Bac-to-Bac system (Invitrogen) and transfected into Sf9 insect cells to produce baculovirus stocks. For protein expression, High Five insect cells at a density of 2 x 106 cells/mL were co-infected with MRT-CRBN-B11 and MRT-DDB1-B05 baculoviruses at a multiplicity of infection (MOI) of 10 and 5, respectively, in the presence of 50 μM ZnAc. Cells were grown at 27°C for 48 hours with shaking before harvesting. Cells were lysed by homogenization in lysis buffer (50 mM Tris pH 8.0, 750 mM NaCl, 1 mM TCEP, 5% glycerol) supplemented with PMSF, EDTA-free protease inhibitors, and benzonase. The lysate was clarified by centrifugation and the MRT-CRBN-B11/DDB1-B05 complex was captured by immobilized metal affinity chromatography using Ni-NTA resin (Qiagen) and eluted with 300 mM imidazole. The eluate was diluted 10-fold and applied to a 5 mL HiTrap Q HP column (GE Healthcare) for ion exchange chromatography using a 50 mM HEPES pH 7.0, 5-100% NaCl gradient. Peak fractions were pooled, concentrated, and further purified by size exclusion chromatography on a Superdex 200 16/600 column (GE Healthcare) pre-equilibrated in 10 mM HEPES pH 7.0, 240 mM NaCl, 1 mM TCEP. The final purified MRT-CRBN-B11/DDB1-B05 complex was concentrated to 38.5 mg/mL, flash frozen in liquid nitrogen, and stored at −80°C.

#### 4. Expression and Purification of VAV1(K782-845C)

pET21b-GST-TEV-Vav1(K782-845C) was transformed into E. Coli BL21 CodonPlus (DE3) cells. These transformed cells were cultured on LB agar plates containing Kanamycin and Ampicillin at 37°C overnight. A single colony was then inoculated into 100 ml of 2YT medium and grown overnight at 37°C. This culture was subsequently scaled up to 2 L of 2YT medium and grown at 37°C until the OD600 reached 0.8. Protein expression was induced with 0.5 mM IPTG at 25°C for 18 hours. The cells were then harvested by centrifugation, yielding 35.5 grams of biomass from the 6 L culture. The biomass was lysed in lysis buffer (20 mM HEPES pH 7.0, 300 mM NaCl, 0.25 mM TCEP, 1 mM PMSF, 5% Glycerol, protease inhibitor, Benzonase) using a homogenizer. The lysate was clarified by centrifugation at 20,000 g for 40 minutes. The supernatant was then loaded onto a Glutathione Sepharose TM 4 Fast Flow column. After washing the column with binding buffer, the protein was eluted with elution buffer (20 mM HEPES pH 7.0, 300 mM NaCl, 0.25 mM TCEP, 5% Glycerol, 10 mM Reduced Glutathione). Further purification was achieved through size-exclusion chromatography (SEC). The concentrated sample was loaded onto a Superdex 75 16/600 pg column equilibrated with SEC buffer (20 mM HEPES pH 7.0, 150 mM NaCl, 0.25 mM TCEP). Fractions were collected and pooled. The protein was stored in a buffer containing 20 mM HEPES pH 7.0, 150 mM NaCl, 0.25 mM TCEP, and 5% Glycerol, with aliquots flash-frozen in liquid nitrogen and stored at −80°C.

For crystallization, a DNA construct encoding a human VAV1 C-terminal SH3 domain fragment (782-845 aa) was inserted into pET21b and expressed as an N-terminal His-tagged fusion in *Escherichia coli* BL21 (DE3) Rosetta cells. Cells were cultured in Terrific Broth, induced with 1 mM IPTG, harvested, and lysed in buffer (20 mM Tris-HCl pH 8.0, 500 mM NaCl, 0.25 mM TCEP, 10% glycerol) supplemented with EDTA-free protease inhibitors (Roche) and 250 U/L Benzonase (Merck). VAV1 was purified by nickel IMAC, dialyzed, treated with TEV protease to remove the His-tag, and further purified by subtractive nickel IMAC followed by size exclusion chromatography (Superdex 75 16/600) in 20 mM HEPES pH 7.0, 100 mM NaCl, 0.5 mM TCEP. The final protein was concentrated to 3.4 mg/mL for crystallization.

#### 5. Expression and Purification of VAV Mutants

The following protein constructs were expressed and purified:

- MRT-Vav1-E20: pET21b with GST-TEV-(Vav1 SH3 domain amino acids K782-845 with mutations R798M, S799R)
- MRT-Vav2SH3-E01: pET21b with GST-TEV-(Vav2 SH3 domain R816-878Q)
- MRT-Vav3 SH3-E01: pET21b-GST-GGG-TEV-Vav3(K788-847E)
- MRT-Vav3 SH3-E02: pET21b-GST-GGG-TEV-Vav3(K788-847E, Y818L, T819N, M821K, S822G, A823Q, N824Q)
- MRT-Vav3 SH3-E03: pET21b-GST-GGG-TEV-Vav3(K788-847E, M804R, R805S, Y818L, T819N, M821K, S822G, A823Q, N824Q)

The proteins were expressed in E. coli BL21 Star (DE3) cells using TB autoinduction medium. The TB medium was prepared with tryptone, yeast extract, glycerol, potassium phosphate salts, glucose, and trace elements solution. Cells were grown at 37°C until OD600 reached 2.0-3.0, then cooled to induction temperatures of 16°C or 25°C and induced with 0.5 mM IPTG. For some constructs, 20 μM ZnSO4 was also added at induction. Cells were harvested after 18-20 hours of expression. The proteins were purified by affinity chromatography using a GST affinity resin. Cell pellets were resuspended in lysis buffer (20 mM HEPES pH 7-7.5, 1 M NaCl, 1 mM TCEP, 1 mM PMSF, EDTA-free protease inhibitors, and Benzonase) and lysed by sonication. Clarified lysates were loaded onto the GST resin and washed with binding buffer (20 mM HEPES pH 7-7.5, 500 mM NaCl, 1 mM TCEP). Proteins were eluted with elution buffer (20 mM HEPES pH 7-7.5, 200 mM NaCl, 1 mM TCEP, 10 mM reduced glutathione). Size-exclusion chromatography was performed as a final polishing step using a Superdex 200 column equilibrated with SEC buffer (20 mM HEPES pH 7-7.5, 200 mM NaCl, 0.5-1 mM TCEP). Purified proteins were concentrated, flash-frozen in liquid nitrogen, and stored at −80°C.

#### 6. Expression and Purification of mTOR-FRB(R2018-2114Q)

The MRT-MTOR-E02 protein was expressed in *Escherichia coli* BL21 Star (DE3) cells using the pET21b vector with an N-terminal His-GST tag followed by a TEV protease cleavage site and coding regions R2018-2114Q. A pre-culture was grown overnight at 37°C in TB medium containing ampicillin. The main culture was inoculated with the pre-culture and incubated at 37°C until the OD600 reached 2.0-3.0 and induced using IPTG. The cell biomass was resuspended in lysis buffer (25 mM Tris-HCl, pH 8.0, 1 M NaCl, 1 mM TCEP, 10% glycerol, 1 mM PMSF, and cOmplete™ EDTA-free protease inhibitor). Cells were lysed using a high-pressure homogenizer. The lysate was clarified by centrifugation. The supernatant was incubated with Glutathione Sepharose 4 Fast Flow resin (GE) at 4°C, and the resin was packed into a gravity column. The column was washed with binding buffer (25 mM Tris-HCl, pH 8.0, 500 mM NaCl, 1 mM TCEP, 10% glycerol) and eluted with elution buffer (25 mM Tris-HCl, pH 8.0, 200 mM NaCl, 1 mM TCEP, 10% glycerol, 10 mM reduced glutathione). His-tagged TEV protease was added to the eluted protein at an enzyme:protein ratio of 1:20 w/w and incubated at 4°C overnight. After cleavage, the sample was loaded onto a gravity column containing Ni-NTA resin pre-equilibrated with reverse binding buffer (25 mM Tris-HCl, pH 8.0, 200 mM NaCl, 1 mM TCEP, 10% glycerol). The column was washed with reverse binding and wash buffers. The protein was eluted using reverse elution buffer (25 mM Tris-HCl, pH 8.0, 200 mM NaCl, 1 mM TCEP, 10% glycerol, 10 mM reduced glutathione, 300 mM imidazole). The pooled fractions from the Ni-NTA column were concentrated using an Amicon Ultra centrifugal filter (10 kDa MWCO), and any precipitates were removed by centrifugation. The sample was loaded onto a Superdex 75 16/600 PG column equilibrated with SEC buffer (25 mM Tris-HCl, pH 8.0, 200 mM NaCl, 1 mM TCEP). The appropriate fractions were pooled and concentrated. The protein was aliquoted, snap-frozen in liquid nitrogen, and stored at −80°C.

#### 7. Expression and Purification of human NEK7

The protein sequence for NEK7, as expressed, includes the N-terminal sequence of NEK7 (M1-S302) followed by a His6-tag for purification. Expression of the protein was carried out in *Escherichia coli* BL21 Star (DE3) cells. Cultures were grown in TB medium supplemented with 50 mg/L ampicillin and 50 mg/L kanamycin. A scale-up culture was inoculated at a 1:10 ratio from an overnight starter culture. The temperature was initially set at 37°C until the optical density (OD600) reached between 2.0 and 3.0. Induction of protein expression was initiated by adding 0.5 mM IPTG, followed by incubation at 16°C for 17 hours. Cell biomass was collected from the culture for subsequent purification. Cells were lysed using a homogenizer at 400 bar for two passes followed by six passes at 900 bar. The lysate was clarified by centrifugation at 13,000 x g for 40 minutes. The clear lysate was incubated with 3 ml of Ni-NTA resin for 1.5 hours. The resin was then packed into a gravity-flow column and washed with 80 ml of washing buffer (20 mM HEPES, pH 7.5, 500 mM NaCl, 1 mM TCEP, 5% glycerol, 15 mM imidazole). The protein was eluted in 25 ml of elution buffer containing 400 mM imidazole. The Ni-IMAC-purified protein was concentrated using an Amicon Ultra centrifugal filter with a 30 kDa molecular weight cutoff (MWCO). Size exclusion chromatography was performed using a Superdex 200 16/600 PG column equilibrated with SEC buffer (20 mM HEPES, pH 7.5, 200 mM NaCl, 1 mM TCEP, 5% glycerol) at a flow rate of 1 ml/min. Fractions were pooled and concentrated, and aliquots were snap-frozen in liquid nitrogen and stored at −80°C. Aliquots were stored in 20 mM HEPES, pH 7.5, 200 mM NaCl, 1 mM TCEP, and 5% glycerol at a concentration of 13.83 mg/ml.

### Crystallographic methods

#### 1. Crystallization of the DDB1-CRBN-mTOR-FRB ternary complex

The protein solution for crystallization was prepared by mixing purified tag-on MRT-CRBN-B02/DDB1-B06 (in 10mM Hepes pH 7.4, 150 mM NaCl, 1 mM TCEP) with FPFT-2216. After incubation for 30 minutes at 4 °C, MRT-mTOR-FRB-E02 (50 mM Tris-HCl pH8.0, 200 mM NaCl, 1 mM TCEP) was added and a final concentration of 80 μM for each component was adjusted, followed by a 1 h incubation at 4 °C and a centrifugation step at 12,000 g for 5 minutes at 4 °C to clear the supernatant. Small crystals were obtained in refined conditions originating from the NeXtal JCSG Core Suite I, containing 70 mM MES pH 6.0, 2.5-4.0 % (w/v) PEG 3,000, and 18-23.5 % (w/v) PEG 200. Seed stocks were prepared from initial crystals and included in subsequent crystallization setups. Final crystals were grown in reservoir solutions containing 70 mM MES pH 6.0, 1.8-3.2 % (w/v) PEG 3,000, and 13-24 % (w/v) PEG 200. For cryoprotection, crystals were transferred into reservoir solution supplemented with 30 % PEG 200 and flash cooled in liquid nitrogen.

#### 2. Crystallization of the DDB1-CRBN-NEK7 ternary complex

The protein solution for crystallization was prepared by mixing purified tag-off MRT-CRBN-B02/DDB1-B06 (in 10 mM Hepes pH 7.4, 150 mM NaCl, 1 mM TCEP) with MRT-3486. After incubation for 30 minutes at 4°C, MRT-NEK7-E06 (50 mM Tris-HCl pH 8.0, 200 mM NaCl, 1 mM TCEP, 5 % (w/v) glycerol) and ADP were added and a final concentration of 70 μM CRBN/DDB1, 80 μM MRT-3486, 70 μM NEK7 and 70 μM ADP was adjusted. After an additional 1 h incubation at 4 °C, the protein solution was centrifuged at 12,000 g for 5 minutes at 4 °C to clear the supernatant. Initial crystal clusters were obtained in a Morpheus I condition containing 100 mM Tris/Bicine pH 8.5, 10 % (v/v) PEG 8,000, 20 % (v/v) ethylene glycol, 30 mM MgCl_2_ and 30 mM CaCl_2_. Crystal clusters were used for seed stock preparation and included in subsequent crystallization setups. Final crystals were obtained by mixing tag-off MRT-CRBN-B02/DDB1-B06 and MRT-NEK7-E06 at equimolar ratios (70 μM final concentration), and 1 mM ADP and 2 mM MgCl_2_ were added before the addition of 100 μM MRT-3486. The sample was centrifuged at 12,000 g for 3 minutes at 4 °C before usage. The protein solution was equilibrated against a reservoir solutions containing 0.029 M HEPES salt, 0.071 M MOPS acid, 0.06 M NaNO3, 0.06 M Na2HPO4, 0.06 M (NH4)2SO4, 11% (w/v) PEG 8,000 and 25 % (v/v) ethylene glycol. Crystals of desired morphology were obtained after multiple rounds of seeding. Crystals were harvested without any further cryoprotection and flash cooled in liquid nitrogen.

#### 3. Crystallization of the DDB1-CRBN-VAV1 ternary complex

CRBN/DDB1-MRT23227-VAV1-SH3c was screened at a ratio of 80 μM : 80 μM : 90 μM. Initial crystal hits were from BCS (Basic Chemical Space) screen D8 condition. Data were collected from crystals grown in 2.6 % PEG Smear Low, 6.1 % PEG Smear Medium, 4.3 % PEG Smear High, 5 % glycerol, 0.1 M CaCl_2_, and 0.1 M MES pH 5.8. Crystals were cryopreserved with 25 % PEG 550 MME and collected at CLS CMCF-ID.

#### 4. Stucture determination

Diffraction data were processed with DIALS (*62*), AIMLESS (*63*) from the CCP4 suite and STARANISO (*64*), providing a primary set of structure factor amplitudes. An alternate set of structure factor amplitudes was generated from DIALS intensity estimates using CARELESS (*65*), for comparative refinement. Initial phases were calculated molecular replacement in PHASER (*66*) using search models derived from PDB entries. After initial refinement with autoBUSTER (*67*) with target model restraints generated from the input model, MR solutions were rebuilt interactively in *COOT* (*68*). Model rebuilding was iterated with refinement variously autoBUSTER or *PHENIX* (*69*) against the primary or alternate set of structure factor amplitudes. Target models for refinement were prepared from template protomers extracted from the current refinement working model with *ALPHAFOLD2* (*70*) as implemented in the local version of COLABFOLD (*71*). Target model restraints were discarded for the concluding refinement iterations. Compound restraints were generated with *GRADE* (*72*). Final models were produced by refinement with PHENIX against the primary structure factor amplitudes with TLS parameterization of the atom displacement parameters with groups defined by *phenix.find_tls.* Data and refinement statistics are in Table S1.

